# Unfolding Suppression: statistical learning drives suppression through dynamic attentional states modulated by threat

**DOI:** 10.1101/2025.09.12.675774

**Authors:** Jingqing Nian, Di Zhang, Yu Zhang, Yu Luo

## Abstract

How does statistical learning drive distractor suppression? A central debate in current research concerns whether the underlying mechanism operates proactively or reactively. Here, we propose a *Temporal Dynamics of Attentional Suppression* (TDAS) hypothesis, successfully reconciling these seemingly contradictory views. Across four visual search experiments, participants faced a probabilistic electric shock when a neutral distractor (Experiments 1a, 1b) or a threat-associated distractor (Experiments 2a, 2b) appeared at a threat-related high-probability location. Trial-averaged results showed faster responses and reduced fixations on distractors, but slower responses and fewer fixations on targets at these locations, consistent with learned distractor suppression. SMART analyses for eye-tracking data further revealed that learned suppression emerged from temporal state transitions: from distractor dominance via attentional limbo states to proactive suppression. Critically, spatial probability gates suppression pathways. At low-probability locations, suppression transitions to limbo states, while at high-probability locations, suppression either unfolds through the full three-state sequence or shifts directly from attentional limbo states to proactive suppression. Furthermore, threat signals accelerated these transitions compared to neutral contexts. Overall, these findings contribute to resolving the ongoing debate about the mechanisms of learned distractor suppression. They also offer a new theoretical perspective by establishing spatiotemporally distributed suppression as a core principle of attentional selection, thereby providing a unifying framework for understanding the dynamic nature of attentional control.

## Introduction

In daily life, we are often distracted by large amounts of information that are not related to our current goals. Given the limits of cognitive resources, it is crucial to understand how the brain predicts and processes distracting information to minimizing its impact and enhancing attentional control. A cognitive process that supports this ability is visual statistical learning, the ability to extract temporal and/or spatial regularities from the environment^1^. Recent research has highlighted the crucial role of visual statistical learning underlying target selection^2–4^ and distractor suppression^5–7^. For instance, Wang and Theeuwes (2018b) demonstrated that participants respond more faster when distractors are more likely to appear in specific locations. This finding indicated that individuals develop a learned distractor suppression at high-probability locations. Such learned distractor suppression is not restricted to the spatial domain, as many studies have demonstrated that distractor interference can be attenuated(Allenmark et al., 2022; Failing & Theeuwes, 2020; Feldmann-Wüstefeld et al., 2021; Ferrante et al., 2018; Goschy et al., 2014; Huang et al., 2021, 2023; Sauter et al., 2021; van Moorselaar et al., 2020, 2021; Zhang et al., 2022; Wang & Theeuwes, 2018a, 2018b, 2018c), or even eliminated, through learning about probable distractor features^18–22^. The learned distractor suppression helps to optimize attentional allocation by reducing interference from irrelevant stimuli.

## 1. The debates on mechanisms of learned distractor suppression

Despite extensive evidence showing that learned distractor suppression reduces interference, the underlying mechanisms remain debated. On the one hand, some studies have shown that learned distractor suppression is initiated before the onset of the search display, indicating proactive suppression^2,12,13,23–29^. For instance, Huang et al. (2022) found that response was faster in the search task when a salient but task-irrelevant color singleton frequently appeared at a high-probability location; and probe detection was slower when the offset dot appeared at this location in the probe task. On the other hand, other studies suggest that learned distractor suppression is an reactive process, involving rapid disengagement of attention from the distractor after initial capture^14,30–33^. For example, Sauter et al. (2021) reported that fixation dwell times were significantly shorter for distractors at high-probability than at low-probability locations. This suggests that attention disengaged more rapidly from high-probability distractor locations, consistent with a reactive mechanism in which suppression is coupled with rapid oculomotor disengagement.

These inconsistencies may occur because the results rely heavily on trials-averaged methods (i.e., aggregate data across trials). Although useful for capturing stable behavioral performance, this approach obscures the trial-by-trial temporal dynamics of attentional control and the interplay between proactive and reactive suppression mechanisms^34^. For instance, reduced distractor interference is often attributed to proactive suppression, yet similar effects could arise from rapid reactive processes. Individuals may differ in their reliance on each mechanism, either trial-by-trial^35,36^ or as a stable trait^37,38^. Furthermore, reactive suppression may compensate when proactive efforts fail. Such variability is masked by averaging but underscores the fluid interplay between these suppression modes.

## 2. Temporal dynamics as a solution to mechanistic debates in learned distractor suppression

Moreover, recent studies suggest that attention operates as a process of accumulating evidence over time^39–43^. As highlighted by Liesefeld et al. (2024), suppression may occur at three distinct temporal stages: (a) before distractor onset, reflecting anticipatory preparation; (b) after distractor onset but before attentional capture, indicating rapid filtering; and (c) after attentional capture. Therefore, it is highly possible that within a given experiment, individuals suppress distractors in high-probability locations through distinct mechanisms. As the experiment unfolds, proactive and reactive suppression may engage at different time points^44^.

Recent years have witnessed a growing adoption of the Smoothing Method for the Analysis of Response Time Courses (SMART) to overcome the limitation of trial-average approach. SMART is a computational model that reconstructs continuous temporal profiles from single-sample-per-trial data^45^. This method facilitates robust statistical inference and has demonstrated considerable utility in eye-tracking research^12,42,43,46–49^. The human oculomotor system offers a valuable window into the attentional process. Specifically, the probability of where a first fixation lands is a sensitive metric reflecting the initial allocation of attentional resources^31,50,51^. Applying SMART to analyze first-fixation landing probabilities, van Heusden et al., (2022) delineated three distinct manifestations of distractor suppression: (1) reactive suppression, following short-lantency saccades, first fixations were more likely to land on distractors than on targets; (2) proactive suppression, following long-latency saccades, the pattern reversed, with first fixations more likely to land on targets; (3) attentional limbo, at intermediate latencies, no reliable difference was observed between the probabilities of landing on distractors versus targets.

However, as learned distractor suppression is a process that accumulates evidence across trials^39–41^, it remains an open question whether these distinct temporal patterns will be evident at a trial-by-trial level. To address this issue, we propose the Temporal Dynamics of Attentional Suppression (TDAS) hypothesis, which posits that learned distractor suppression is not a unitary process but rather emerges from the dynamic integration of distinct suppression mechanisms within the priority map over time, transitioning through reactive suppression, an attentional limbo state, and proactive suppression. Accordingly, the current study employs SMART to examine how attentional mechanisms dynamically shift across individual trials during the development of learned distractor suppression.

## 3. Distractor salience as a modulator of learned distractor suppression

Furthermore, distractor salience represents a critical factor modulating the development and strength of learned suppression. Previous studies have demonstrated that physical salience of distractors (e.g., color, luminance, and size) can potentiate learned distractor suppression at high-probability locations (Failing & Theeuwes, 2020; Gong & Theeuwes, 2021). For instance, Gong and Theeuwes (2021) manipulated the salience and spatial distribution probability of distractors within an additional singleton paradigm. Their results showed that highly salient distractors at high-probability locations were suppressed more strongly than their low-salience counterparts at the same locations, suggesting that the visual system implements salience-dependent suppression through statistical learning. In contrast to physical salience, the role of threat-related salience in facilitating learned distractor suppression remains unclear. Threat-related stimuli, due to their evolutionary significance, are often prioritized and more resistant to suppression ^52–59^.

However, emerging evidence indicates that individuals can, under certain conditions, learn to suppress threat-related stimuli^60–62^. The nature of this suppression appears contentious. For example, Theeuwes et al. (2025) reported that threat-related distractors at high-probability locations were effectively suppressed, but no stronger than neutral distractors. Conversely, Nian et al. (2025) observed a stronger suppression effect for threat stimuli. This discrepancy may arise because the learned suppression of threat-related distractors involves a dynamic integration of multiple mechanisms across different temporal stages, resulting in a more complex time course. Notably, these findings were primarily based on trial-averaged data, which obscures moment-to-moment dynamics. Consequently, investigating how threat-related stimuli influence the trial-by-trial dynamics of learned distractor suppression is crucial. Such an approach is essential for elucidating how motivational salience interacts with statistical learning to shape attentional control over time.

To this end, we conducted four experiments examining whether learned distractor suppression involves a dynamic integration of proactive and reactive mechanisms over time, and whether this process is modulated by threat-related salience. We employed a modified additional singleton task featuring two high-probability distractor locations. In Experiments 1a and 1b, we tested how the threat associated with a spatial location influences learned suppression by pairing one of the high-probability locations with an aversive electric shock (e.g., the shock paired location, CsPH). In Experiments 2a and 2b, we extended this approach to examine how a threat linked to a specific feature-location conjunction modulates learned suppression. This was achieved by pairing the shock with the specific combination of the distractor’s feature and its high-probability location (e.g., a specific color at a specific location, CsP–CsPH). To directly capture the temporal dynamics of attentional location, we employed eye tracking in Experiments 1b and 2b. We analyzed first-fixation landing probabilities using the SMART model, which reconstructs continuous temporal profiles from single-trial data and reveals the temporal dynamics of suppression mechanisms.

## Experiment 1a

Experiment 1a tested whether learned suppression of distractors would be stronger when they appeared at a high-probability location that was threat-associated than at another one that was not associated with threat. The primary hypothesis was that participants would develop learned suppression for high-probability locations, and the threat-associated salience would modulate the learned suppression. Specifically, if participants acquired suppression, we expected shorter response times (RTs) for the high-probability locations than the low-probability locations. Furthermore, if threat-associated salience potentiates learned suppression, we expected shorter RTs in the threat-associated salience high-probability location than the non-threat-associated location.

## Materials and Methods

### Participants

The sample size was determined based on the method outlined in a previous study (Le Pelley et al.,2022). A priori power analysis using G*Power (version 3.1; Faul et al., 2007) indicated that a sample size of 29 would provide over 80% power to detect a medium within-subject effect size (*d* = 0.54) at an alpha level of 0.05.

A total of 32 participants (26 females; age range 19-25 years) participated in Experiment 1a. All participants had normal or corrected-to-normal vision, normal color vision, and were right-handed. The participants provided written informed consent approved by the Committee on Human Research Protection of the School of Psychology, Guizhou Normal University (GZNUPSY.NO 2022 E [007]). Participants were compensated at a rate of ¥30 per hour upon completing the study.

### Stimuli and Apparatus

The stimuli consisted of eight shapes (1.8° × 1.8°), including one diamond and seven circles. Inside each shape was a black (RGB: 5, 5, 5) oriented bar (0.8° × 0.08°) in two possible directions (vertical or horizontal). In the distractor-present condition, one circle was filled with either orange (RGB: 255, 69, 0) or blue (RGB: 0, 128, 255). Each participant was assigned only one of these colors. In all other conditions, all shapes were filled with gray (RGB: 109, 109, 109).

Screen luminance and color were measured and calibrated using the PsyCalibrator toolkit and SpyderX^63^. The calibrated parameters were then applied during the stimulus presentation phase of the experiment. All stimuli were generated using Psychtoolbox 3.0 running in MATLAB (version 2020b) and were presented on a dark gray background (RGB: 65, 65, 65). The stimuli were displayed on a 21.5-inch LCD monitor with a resolution of 1920 × 1080 and a refresh rate of 60 Hz, and the viewing distance was 72 cm.

### Experimental Tasks

#### Shock work-up task

A multi-channel electrical stimulator (Type: SXC-4A, Sanxia Technique Inc., Beijing, China) was attached to the participant’s left wrist and delivered stimulation through a pair of Ag/AgCl surface electrodes. The shock intensity was calibrated for each participant individually to reach a level that was rated as highly uncomfortable, yet not painful. Specifically, it was set to a score of 8 on a subjective discomfort scale ranging from 0 (no sensation) to 10 (extremely painful). The final intensity used in the visual search task was calculated as the average of three shocks, each rated as 8 on the 11-point scale.

#### Visual search Task

As shown in Figure 1A, each trial began with a central fixation for a random duration between 500 and 800ms. Then, a search display consisting of one diamond (the target shape) and seven circles (the non-target shape) presented. The target was defined by a horizontal or vertical bar embedded within a diamond. Participants were instructed to indicate the orientation of the target bar by pressing a key, the key pressing was counterbalanced across participants. The search display remained on the screen for a maximum of 1500 ms. Whether or not the participant responded within this time, the array then disappeared. If the response was incorrect or no response was made within 1500 ms, an auditory beep was presented as a feedback.

**Figure 1.**
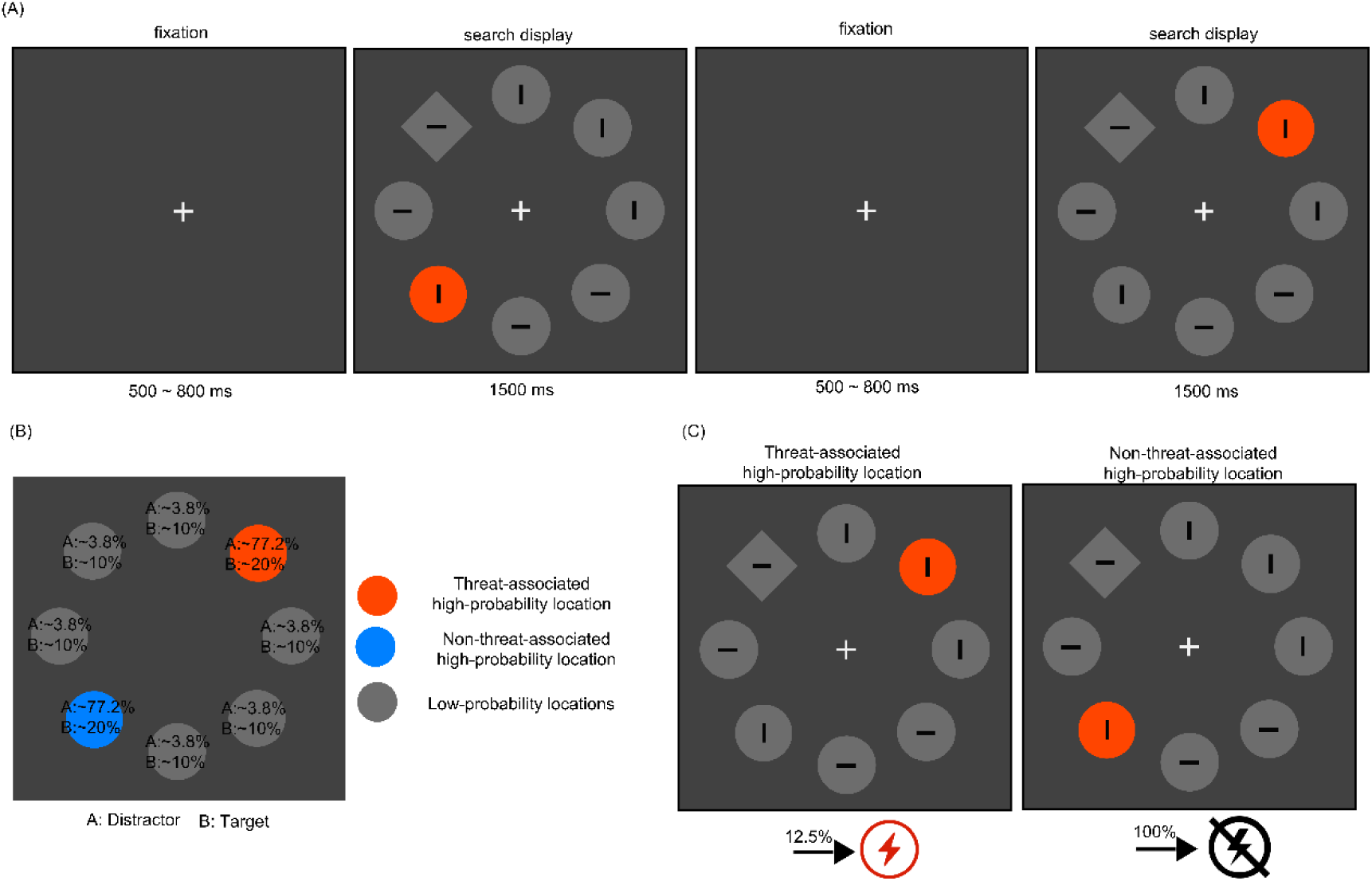
Tasks and Design for Experiment 1a *Note.* Panel A: Trial sequence of the visual search task. Participants were instructed to report the orientation of the bar inside the diamond. In the distractor-present condition, one of the non-target circles was replaced with a singleton of a different color. Panel B: An example of the spatial and salience regularities of the distractor. The two high-probability distractor locations are shown in orange and blue, and the low-probability locations are shown in gray. The percentages at each location indicate the probability that a distractor category or the target presented at that specific location. Panel C: Overview of the threat conditioning trial. The trial includes two conditions: when the distractor was presented at the threat-associated high-probability location (CsPH), 12.5% of those trials are paired with an electric shock. In contrast, when the distractor was presented at a non-threat-associated high-probability location (CsMH) or at any of the low-probability locations (LPL), it was never paired with a shock. See the online article for the color version of this figure.

Each participant completed eight blocks of the visual search task. Each block consisted of 124 trials: 104 distractor-present trials and 20 distractor-absent trials. In each block, two opposite positions out of eight were randomly selected as threat-associated high-probability location and non-threat-associated high-probability location, respectively. The remaining six positions served as low-probability distractor locations. These positions were assigned counterbalanced across participants (see Figure 1B).

In the distractor-present condition, three types of trials were included: (1) the distractor presented at threat-associated high-probability location (CsPH, *trial number* = 40); (2) the distractor presented at non-threat-associated high-probability location (CsMH, *trial number* = 40); (3) the distractor presented at low-probability location (LPL, *trial number* = 24, with each of six locations presented four times). Notably, participants were at risk of receiving a shock only during CsPH trials. Specifically, in each block, five out of 40 CsPH trials were randomly selected to deliver an electric shock, yielding a probability of 12.5% (see Figure 1C). The shock delivery occurred at search array onset and was determined solely by the stimulus location, independent of participants’ responses. To prevent potential confounding effects, trials involving shocks were excluded from analysis. In the distractor-absent condition, the target was presented two times at each of the six low-probability locations, and four times at each of the two high-probability locations.

Within each block, the order of trials and target positions was randomized. After each block, participants received feedback on their average response time and accuracy. The feedback reminded participants of their prior block performance and encouraged them to respond as quickly and accurately as possible in the next block.

#### Implicit learning assessment

After completing the visual search task, participants answered a series of questions to assess their awareness of the spatial distribution of the distractors. First, they were asked if they had noticed any spatial imbalance in the occurrence of the distractor. Then, they were asked to identify the location with the highest probability of a distractor for both the threat- and non-threat-associated location.

## Data Analysis

Behavioral data analyses were conducted in R using RStudio. Accuracy (ACC) was examined using generalized linear mixed-effects models (GLMMs), while response times (RT) were examined using linear mixed-effects models (LMMs). These models were implemented using the *lme4* and *lmerTest* packages ^64^.

Before analysis, the response time (RT) data were preprocessed as follows: (1) Trials with shock delivering were excluded; (2) The first two trials of each block were excluded; (3) trials with incorrect responses were removed; (4) trials with RTs shorter than 200 ms were excluded; (5) trials were excluded if a participant’s reaction time in a given condition deviated by more than ±2.5 standard deviations from their mean response time in that condition; (6) trials involving shocks were excluded from analysis; and (7) participants with fewer than 70% of the total trials after preprocessing were excluded from further analysis. After preprocessing, the proportion of excluded trials ranged from 10.58% to 27.02%, with an average of 16.06% ± 0.75%.

To examine whether individuals exhibited learned suppression at high-probability locations, a mixed-effects linear model was fitted separately to the distractor-absent and distractor-present conditions. In the distractor-absent condition, both ACC and RT were modeled with stimulus location (LocP) as a fixed effect and subject as a random effect (model: ACC/RT ∼ LocP + (1 | Subject)). In the distractor-present condition, distractor location included as fixed effects with subject and target-distractor distance (TDD) as random effects (model: ACC/RT ∼ LocP + (1 | subject) + (1 | TDD)). Post-hoc comparisons were conducted using the *emmeans* package with Tukey adjustments for multiple comparisons.

## Transparency and Openness

All data, analysis scripts, and task codes will be made publicly available via the *Science Data Bank* upon acceptance of the manuscript. This study was not pre-registered.

## Results

### Distractor-absent condition

As shown in Figure 2, accuracy analysis revealed that the main effect of distractor location was not significant (*χ*²_(2)_ = 1.18, *p* = 0.40).

**Figure 2.**
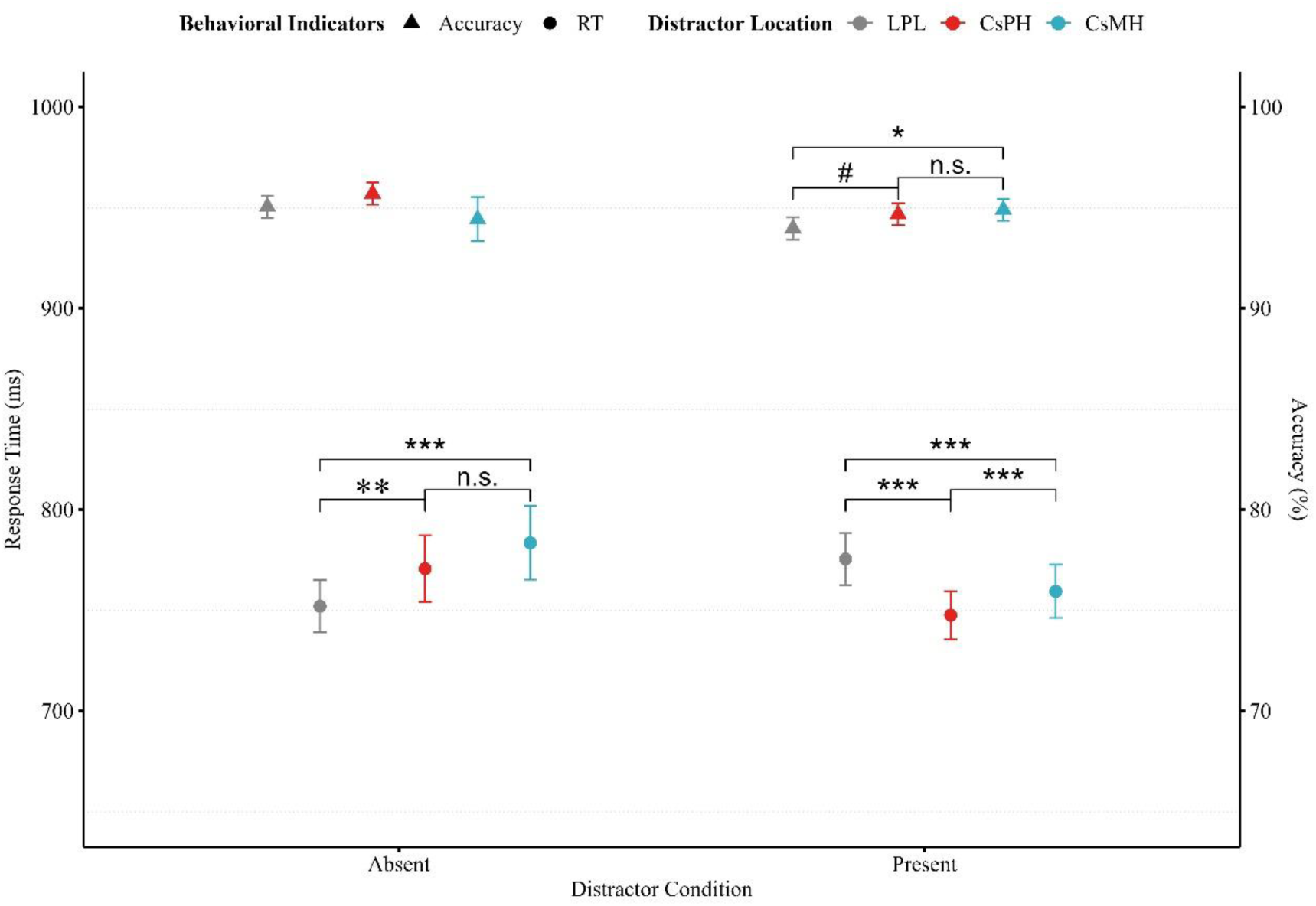
The results of search performance to target across different conditions in Experiment 1a. *Note.* Behavioral results for visual search. The left Y-axis shows response time, represented by solid circles, and the right Y-axis shows accuracy, represented by solid triangles. All error bars represent the within-subject standard error of the mean. LPL: low-probability distractor locations; CsPH: threat-associated high-probability location; CsMH: non-threat-associated high-probability location. *** indicates *p* < 0.001, ** indicates *p* < 0.01, * indicates *p* < 0.05, and n.s. indicates not significant (*p* > 0.05). See the online article for the color version of this figure.

Response time analysis revealed a significant main effect of distractor location, (*χ*²_(2)_ = 26.83, *p* < 0.001). Compared to low-probability locations (752.05 ± 12.95 ms), responses were slower at both threat-associated high-probability location (770.68 ± 16.48 ms, *b* = 18.36, *SE* = 6.38, *t*_(44,445)_ = 2.88, *p* = 0.004, 95% CI [5.85, 30.87]; *β* = 0.04, 95% CI [0.01, 0.07]) and non-threat-associated high-probability location (783.54 ± 18.24 ms, *b* = 31.22, *SE* = 6.38, *t*_(44,445)_ = 4.89, *p* < 0.001, 95% CI [18.72, 43.73]; *β* = 0.07, 95% CI [0.04, 0.10]). However, no significant difference was found between the two high-probability locations (i.e., threat- and non-threat-associated (*b* = 12.86, *SE* = 7.80, *t*_(44,445)_ = 1.65, *p* = 0.10, 95% CI [−2.43, 28.15]; *β* = 0.03, 95% CI [−0.01, 0.06]).

### Distractor-present condition

As shown in Figure 2, accuracy analysis revealed a significant main effect of distractor location (*χ*²_(2)_ = 6.88, *p* = 0.03). Compared to low-probability locations (93.97 ± 0.55%), accuracy were higher at both threat-associated high-probability location (94.67 ± 0.56%, *b* = 0.13, *SE* = 0.07, *z* = 1.92, *p* = 0.054, 95% CI [−0.003, 0.27], *OR* = 1.14) and non-threat-associated high-probability location (94.90 ± 0.54%, *b* = 0.18, *SE* = 0.07, *z* = 2.58, *p* = 0.01, 95% CI [0.04, 0.32], *OR* = 1.20). However, no significant difference was found between the two high-probability locations (*b* = 0.05, *SE* = 0.06, *z* = 0.75, *p* = 0.45, 95% CI [-0.08, 0.17], O*R* = 1.05).

Response time analysis revealed a significant main effect of distractor location (*χ*²_(2)_ = 31.40, *p* < 0.001). Compared to low-probability locations (775.50 ± 13.04 ms), Responses were faster at both threat-associated high-probability location (747.64 ± 11.95 ms, *b* = −9.294, *SE* = 2.931, *t*_(22,128)_ = −3.17, *p* = 0.002, 95% CI [−15.04, −3.55]; *β* = −0.03, 95% CI [−0.04, −0.01]) and non-threat-associated high-probability location (759.44 ± 13.22 ms, *b* = −16.00, *SE* = 2.86, *t*_(22,112)_ = −5.59, *p* < 0.001, 95% CI [−21.60, −10.39]; *β* = −0.04, 95% CI [−0.06, −0.03]). Additionally, Responses were also faster at the threat-associated high-probability location than in the non-threat-associated high-probability location (*b* = −6.65, *SE* = 2.55, *t*_(22,129)_ = −2.60, *p* = 0.009, 95% CI [−11.65, −1.65]; *β* = −0.02, 95% CI [−0.03, −0.004]).

### Awareness of statistical regularities

Ten of the 32 participants reported that they did not notice whether the distractor appeared more or less frequently in a specific location. These participants gave an average confidence rating of 4.00 ± 0.54 on a 7-point likert scale. The other 22 participants reported noticing a spatial imbalance and gave an average confidence rating of 3.77 ± 0.38. However, none of these participants correctly identified the high- or low-probability distractor locations as defined in the experiment. These results suggest that the learned suppression effect was driven by implicit awareness.

## Discussion

In Experiment 1a, the trial-averaged results replicated the classic finding that high-probability locations lead to learned distractor suppression. Participants responded faster when a neutral distractor appeared at a high-probability location than when it appeared at low-probability location. This benefit was further amplified when the high-probability location was associated with a threat. These results support the saliency-specific mechanism of distractor suppression, which posits that stimuli with higher salience are more likely to elicit stronger learned suppression at high-probability locations^9,65^.

However, trial-averaged data obscure within-participant temporal dynamics, making it impossible to determine whether suppression reflects genuine enhancement or resistance over time. Consequently, it remains unclear whether learning trajectories at threat- and non-threat-associated locations unfold similarly. To address this issue, Experiment 1b examined the learning process on a trial-by-trial basis. Using eye tracking and the SMART approach, we reconstructed the time courses of first fixations to distinguish reactive suppression from proactive suppression.

## Experiment 1b

In Experiment 1b, eye-tracking and the SMART method were employed to examine whether the temporal course of learned distractor suppression differed between threat-associated and non-threat-associated high-probability locations. As Liesefeld et al. (2024) have highlighted, suppression may occur at three temporal stages. We hypothesize that learned suppression undergoes a trial-level transition from reactive to proactive control at high-probability locations, and that threat modulates this progression. With reactive suppression, the first fixation is significantly more likely to land on the distractor than the target. In contrast, under proactive suppression, the first fixation is more likely to be directed toward the target. When there is no difference in the likelihood of fixating on either, the outcome is identified as attentional limbo.

## Materials and Methods

### Participants

The sample size calculation method and participants’ recruitment criteria were identical to those used in Experiment 1a. Forty-seven participants (36 females; age rang 19-25 years) participated in the current experiment.

### Stimuli and Apparatus

The stimuli and apparatus used in the current experiment were identical to those used in Experiment 1a.

### Experimental Tasks

**Shock work-up task**: identical to that in Experiment 1a.

**Visual search Task**: The behavioral task was identical to Experiment 1a. In the current experiment, the eye movement data were recorded using an EyeLink 1000 Plus eye tracker during the visual search task. The sampling rate of eye tracking was set to 1000 Hz, and monocular gaze data were recorded from the left eye. A 9-point calibration procedure was used to ensure accuracy, with calibration error maintained within 0.5°. Calibration was repeated before the start of each block. Additionally, at the beginning of each trial, participants were required to maintain their gaze within a 1° radius of the central fixation point for 1000 ms before the visual search display appeared.

**Implicit learning assessment**: identical to that in Experiment 1a.

## Data Analysis

The behavioral data preprocessing procedure was identical to that employed in Experiment 1a. After preprocessing, the proportion of trials removed ranged from 9.88% to 27.12%, with an average of 18.23% ± 0.67%.

The eye-tracking data were analyzed as follows: First, fixations that were less than 1° apart and shorter than 100 ms were merged (Hooge et al., 2022). Subsequently, two concentric virtual circles were drawn on-screen with a radius of 50 pixels and another with a radius of 350 pixels (Figure 3). Each circle was divided into eight equal segments using 45° angles. The region of interest (ROI) was defined as the annular area between the inner and outer circles. Fixations beyond the outer circle were excluded. The first fixation was operationalized as the first gaze position entering a stimulus region after a saccade exited the central fixation circle.

**Figure 3.**
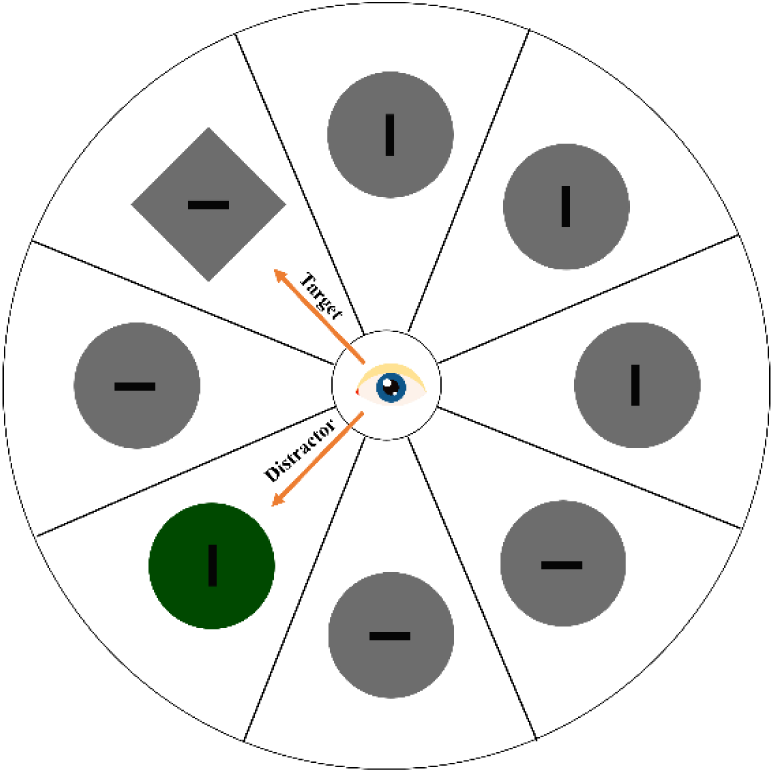
Schematic illustration of the regions of interest (ROIs) *Note.* Two concentric circles (radii: 50 and 350 pixels) were centered on the display, each divided into eight equal angular segments (45° per segment). The region of interest (ROI) for fixation analysis was defined as the annular area between the inner and outer circles. Each numbered sector served as both a distinct ROI and a designated stimulus presentation location in the visual search task. First fixations were classified as on-target if they landed at the target area, on-distractor if they landed at the distractor area, and elsewhere otherwise.

Second, generalized linear mixed-effects models (GLMMs) were used to analyze the probability that participants’ first fixation landing on specific ROI. In the distractor-absent condition, location probability was included as a fixed factor, and the probability that the first fixation landing on the target was used as the dependent variable. Participant variability was modeled by including subject as a random effect. The model was specified as: 𝑃_𝑓𝑖𝑥_𝑇_ ∼ LocP + (1 ∣ Subject). In the distractor-present condition, location probability was included as a fixed factor, with the probability that first fixation landing on the distractor as the dependent variable. Participant variability was accounted for by including subject and target-distractor distance (TDD) as random effects. The model was specified as: 𝑃_𝑓𝑖𝑥_𝐷_ ∼ LocP + (1 ∣ Subject) + (1 ∣ TDD) . Statistically significant differences between conditions and factor levels were assessed via post-hoc pairwise comparisons using the *emmeans* package, with *p*-values adjusted via Tukey’s method for multiple comparisons.

To further explore the attentional mechanisms and temporal dynamics of learned distraction suppression, we used the SMART method to analyze the probability of first-fixation landings^42,45^. First, Gaussian smoothing was applied to the data along the trial axis for each participant. Then, a weighted average was computed to construct a group-level time course. Next, cluster-based permutation testing was used to examine differences between time courses or deviations from a baseline. Specifically, the analysis identified clusters of consecutive time points that exhibited significant differences. For each cluster, a cluster-level *t*-value was calculated by summing the *t*-values within the cluster. This cluster-level statistic was then compared to a threshold determined from randomly permuted datasets. A cluster was considered significant if its *t*-value exceeded the 95th percentile of the permutation distribution.

For all experiments, temporal smoothing was applied to the fixation probability data using a Gaussian kernel with a standard deviation of 30 trials and a step size of one trial. Spatial-temporal clusters were then identified via permutation t-tests with 1000 iterations and cluster-defining threshold α level of 0.05. For non-target and non-distractor items, the average landing probabilities were computed using the SMART method. To analyze the temporal dynamics of visual attention, cluster-based permutation t-tests were used to identify significant differential periods during which the probability of a first fixation landing on a target or on a distractor. If the first fixation was more likely to land on distractors than on targets, this is consistent with the reactive suppression account. If the first fixation landed more often on targets than on distractors, this is consistent with the proactive suppression account. Otherwise, if there was no difference between the first fixation landing on targets and on distractors, this may indicate attentional limbo.

## Results

### Behavioral results

#### Distractor-absent condition

Accuracy analysis revealed a marginally significant main effect of distractor location (*χ*²_(2)_ = 5.47, *p* = 0.06). Accuracy was higher at low-probability location (97.05 ± 0.36%) compared to threat-associated high-probability location (95.94 ± 0.65%, *b* = −0.34, *SE* = 0.16, *z* = −2.15, *p* = 0.031, *OR* = 0.72, 95% CI [−0.64, −0.03]). However, no significant difference between low-probability location and non-threat-associated high-probability location (96.28 ± 0.71%, *b* = −0.25, *SE* = 0.16, *z* = −1.53, *p* = 0.13, OR = 0.78, 95% CI [−0.56, 0.07]). The difference between the two high-probability locations was also not significant (*b* = 0.09, *SE* = 0.19, *z* = 0.48, *p* = 0.63, 95% CI [−0.28, 0.46], *OR* = 1.10).

Response time analysis revealed a significant main effect of distractor location (*χ*²_(2)_ = 28.59, *p* = 0.001). Compared to low-probability location (699.52 ± 12.40 ms), Responses were slower at both threat-associated high-probability location (721.52 ± 16.37 ms, *b* = 23.51, *SE* = 5.01, *t*_(6,434)_ = 4.70, *p* < 0.001, 95% CI [13.70, 33.32], *β* = 0.05, 95% CI [0.03, 0.08]) and non-threat-associated high-probability location (718.09 ± 15.52 ms, *b* = 18.22, *SE* = 5.00, *t*_(6,434)_ = 3.65, *p* < 0.001, 95% CI [8.42, 28.01]; *β* = 0.04, 95% CI [0.02, 0.07]). However, no significant difference emerged between the two high-probability locations (*b* = −5.29, *SE* = 6.12, *t*_(6,434)_ = −0.86, *p* = 0.39, 95% CI [−17.30, 6.72]; *β* = −0.01, 95% CI [−0.04, 0.02]).

#### Distractor-present condition

As shown in Figure 4, accuracy analysis revealed a significant main effect of distractor location (*χ*²_(2)_ = 15.14, *p* = 0.001). Compared to low-probability location (94.84 ± 0.49%), accuracy was higher at both threat-associated high-probability location (95.75 ± 0.37%, *b* = 0.206, *SE* = 0.063, *z* = 3.29, *p* < 0.001, 95% CI [0.08, 0.33], *OR* = 1.23) and non-threat-associated high-probability location (95.83 ± 0.40%, *b* = 0.23, *SE* = 0.06, *z* = 3.60, *p* < 0.001, 95% CI [0.10, 0.35], *OR* = 1.25). However, no significant difference was found between the two high-probability locations (*b* = 0.02, *SE* = 0.06, *z* = 0.35, *p* = 0.73, 95% CI [−0.09, 0.13], *OR* = 1.02).

**Figure 4.**
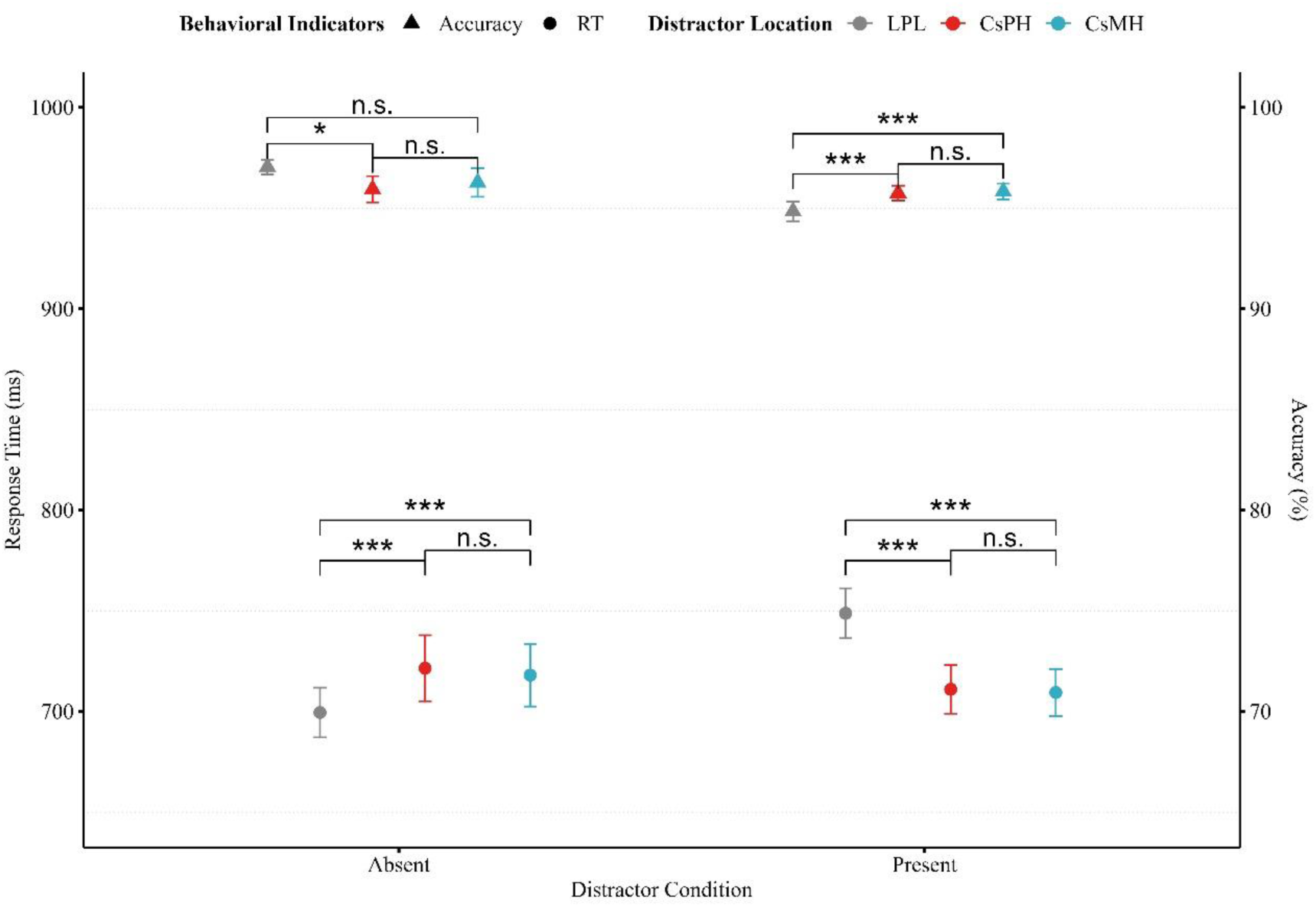
The results of search performance to target across different conditions in Experiment 1b. *Note.* Behavioral results for visual search. The left Y-axis shows response time, represented by solid circles, and the right Y-axis shows accuracy, represented by solid triangles. All error bars represent the within-subject standard error of the mean. LPL: low-probability distractor locations; CsPH: threat-associated high-probability location; CsMH: non-threat-associated high-probability location. *** indicates *p* < 0.001, ** indicates *p* < 0.01, * indicates *p* < 0.05, and n.s. indicates not significant (*p* > 0.05). See the online article for the color version of this figure.

Response time analysis revealed a significant main effect of distractor location, χ²_(2)_ = 335.20, *p* < 0.001. Compared to low-probability location (748.79 ± 12.41 ms), Responses were faster at both threat-associated high-probability location (711.02 ± 12.06 ms, *b* = −37.18, *SE* = 2.35, *t*_(31,590)_ = −15.83, *p* < 0.001, 95% CI [−41.79, −32.58]; *β* = −0.10, 95% CI [−0.11, −0.09]) and non-threat-associated high-probability location (709.45 ± 11.64 ms, *b* = −38.77, *SE* = 2.29, *t*_(31,590)_ = −16.94, *p* < 0.001, 95% CI [−43.26, −34.29]; *β* = −0.11, 95% CI [−0.12, −0.10]). However, no significant difference was found between the two high-probability locations (*b* = −1.59, *SE* = 2.05, *t*_(31,590)_ = −0.78, *p* = 0.44, 95% CI [−5.61, 2.43]; *β* = −0.004, 95% CI [−0.02, 0.01]).

### Results of First Fixation Landing Probability

#### Distractor-absent condition

As shown in Figure 5A, first fixation landing on target analysis revealed a significant main effect of distractor location (*χ*²_(2)_ = 32.66, *p* < 0.001). Compared to low-probability distractor locations (44.73 ± 2.72%), the probability was lower at both threat-associated high-probability location (38.14 ± 3.50%, *b* = −0.31, *SE* = 0.07, *z* = −4.36, *p* < 0.001, 95% CI [−0.45 to −0.17], *OR* =0.74) and non-threat-associated high-probability location (37.68 ± 3.46%, *b* = −0.33, *SE* = 0.07, *z* = −4.63, *p* < 0.001, 95% CI [−0.47, −0.19], *OR* =0.72. However, no significant difference was found between the two high-probability locations (*b* = −0.02, *SE* = 0.09, *z* = −0.22, *p* = 0.83, 95% CI [−0.19, 0.15], *OR* =0.98).

**Figure 5.**
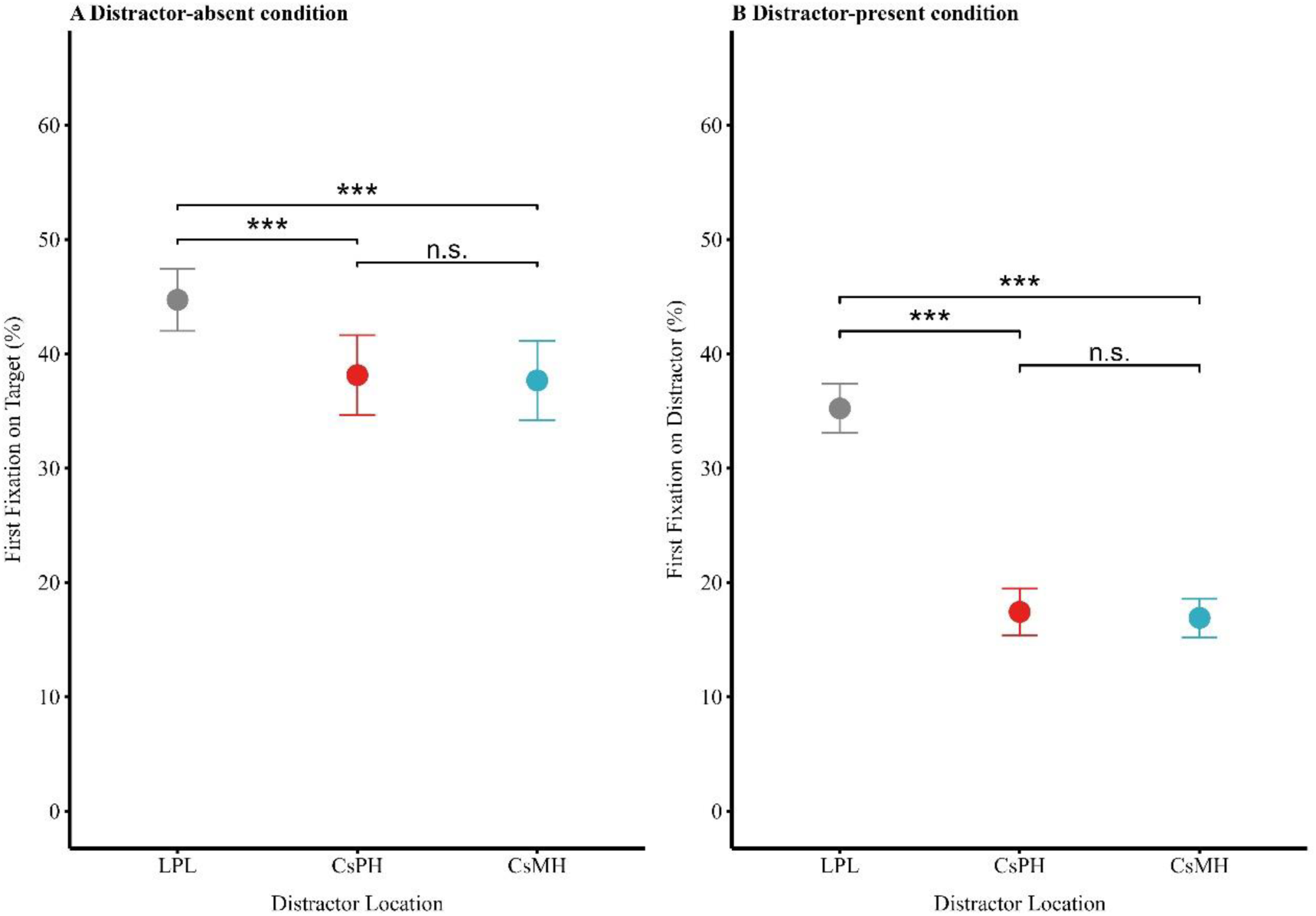
The results of first fixation landing probability across different conditions in Experiment 1b. *Note* Panel A: The probability of the first fixation landing on the target when it appeared at a low-probability location, a threat-associated high-probability location, or a non-threat-associated high-probability location in the distractor-absent condition. Panel B: The probability of the first fixation landing on distractor when the distractor appeared at a low-probability location, a threat-associated high-probability location, or a non-threat-associated high-probability location in the distractor-present condition. All error bars represent the within-subject standard error of the mean. LPL: low-probability distractor locations; CsPH: threat-associated high-probability location; CsMH: non-threat-associated high-probability location. *** indicates *p* < 0.001, ** indicates *p* < 0.01, * indicates *p* < 0.05, and n.s. indicates not significant (*p* > 0.05). See the online article for the color version of this figure.

### Distractor-present condition

As shown in Figure 5B, first fixation landing on the distractor analysis revealed a significant main effect of distractor location (*χ*²_(2)_ = 1123.40, *p* < 0.001). Compared to low-probability location (35.23 ± 2.15%), the probability was lower at both threat-associated high-probability location (17.43 ± 2.02%, *b* = −1.01, *SE* = 0.04, *z* = −28.23, *p* < 0.001, 95% CI [−1.08, −0.94], *OR* =0.36) and non-threat-associated high-probability location (16.91 ± 1.70%, *b* = −1.05, *SE* = 0.04, *z* = 29.88, *p* < 0.001, 95% CI [−1.11, −0.98], *OR* =0.35). However, no significant difference was found between the two high-probability locations (*b* = −0.03, *SE* = 0.04, *z* = −0.88, *p* = 0.38, 95% CI [−0.10, 0.04], *OR* =0.97).

## SMART Results for First Fixation Landing Probability

For distractors appeared at a low-probability location (see Figure 6A), we observed a strong initial attentional capture effect. Specifically, from Trial 1 to 277, the first fixation was significantly more likely to land on the distractor than on the target (*p* < 0.05). However, this capture effect was transient, as no significant difference in first-fixation probability was found between the distractor and the target from Trial 278 onward (*p* > 0.05).

**Figure 6.**
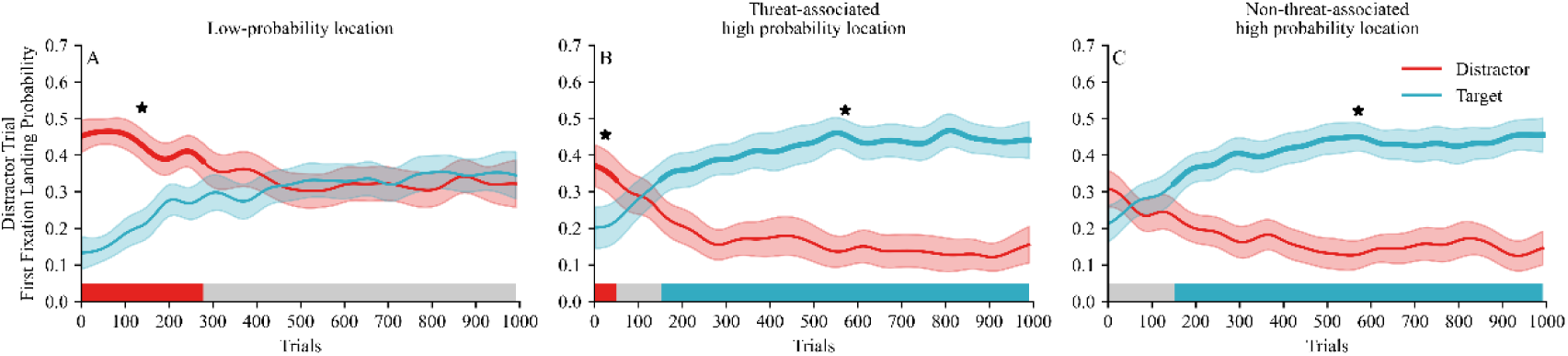
Difference functions reflecting the net distractor and target effects across trials in Experiments 1b *Note.* Panels A–C show data from trials with a distractor present for three conditions: low-probability locations (Panel A), threat-associated high-probability locations (Panel B), and non-threat-associated high-probability locations (Panel C). The shaded areas denote 95% confidence intervals, and the bold lines indicate the time points at which the probability of the first fixation differs significantly between the distractor and the target. * indicates *p* < 0.05. On the x-axis, the red-shaded region represents reactive suppression, the gray region represents attentional limbo, and the blue region represents proactive suppression. See the online article for the color version of this figure.

For distractors appeared at the threat-associated high-probability location (see Figure 6B), we also observed an initial attentional capture effect. Specifically, from Trial 1 to 48, the first fixation was more likely to land on the distractor than on the target (*p* < 0.05). Intriguingly, an attentional limbo phase then emerged between Trials 49 and 151, where no significant difference in first-fixation probability was found between the distractor and the target (*p* > 0.05). Following this, from Trial 152 onward, a pattern of proactive suppression was observed, the first fixation was significantly more likely to land on the target than on the distractor (*p* < 0.05).

For distractors appeared at the non-threat-associated high-probability location (see Figure 6C), we observed a markedly different pattern. During the initial 150 Trials, the first fixation was equally likely to land on the distractor as on the target (*p* > 0.05), indicating an attentional limbo phase. However, beginning with Trial 151, a significant proactive attentional suppression emerged, with the first fixation being significantly more likely to land on the target than on the distractor (*p* < 0.05).

These results illustrate the dynamic evolution of learned distractor suppression, which follows distinct trajectories depending on the salience of location. At low-probability locations, it transitions from reactive suppression to an attentional limbo. At threat-associated high-probability locations, it evolves from reactive suppression, through an attentional limbo, to proactive suppression. At non-threat-associated high-probability locations, it shifts from an attentional limbo to proactive suppression.

## Awareness of statistical regularities

Twenty-three of the 47 participants reported that they did not notice whether the distractor appeared more or less frequently in a specific location. These participants gave an average confidence rating of 4.00 ± 0.31 on a 7-point scale. The remaining 24 participants reported noticing a spatial imbalance and gave an average confidence rating of 4.00 ± 0.29. However, none of these participants correctly identified the high- or low-probability distractor locations as defined in the experiment. These results suggest that the learned suppression effect was driven by implicit awareness.

## Discussion

In Experiment 1b, the trial-averaged results were similar to those of Experiment 1a. Participants responded more quickly when distractors appeared at high-probability locations. However, unlike Experiment 1a, there was no significant difference in response time between distractors at threat-associated high-probability locations versus non-threat-associated high-probability locations. One possible explanation for this null effect is that high-probability locations are subject to "blanket" suppression, which prevents any item from capturing attention, regardless of its threat association.

Although trial-average results suggested that individuals learned to suppress distractors in high-probability locations, SMART analysis revealed that this behavioral benefit does not arise from a unitary attentional control mechanism. Instead, it reflects the dynamic temporal integration of multiple, distinct suppression mechanisms throughout the learning process. Specifically, the patterns of attentional control evolved along distinct trajectories: at low-probability locations, from reactive suppression to a sustained attentional limbo; At non-threat-associated high-probability locations, from attentional limbo toward proactive suppression; At threat-associated high-probability locations, through a more complex progression from reactive suppression, via attentional limbo, to robust proactive suppression. These results indicate that although threat-related locations modulate the trajectory of attentional suppression, they do not diminish the overall behavioral advantage for target processing. Rather, threat signals enhance the attentional priority of their associated locations, thereby accelerating and facilitating the development of higher-order proactive suppression mechanisms.

In summary, whereas traditional trial-averaged approaches support the stability of learned distractor suppression at high-probability locations, the time-resolved SMART analysis demonstrates that this benefit emerges from heterogeneous and dynamic underlying cognitive processes. Critically, in the threatening location, distractor suppression not only emerges earlier, but also transitions more readily from reactive to proactive suppression. These findings provide novel empirical evidence for how threat modulates the dynamic allocation of attentional resources, and underscore the necessity of looking beyond trial-averaged data to understand the rich temporal dynamics of cognitive processes.

## Experiment 2a

Building on the role of threat-associated locations in learned distractor suppression established in Experiment 1, we next sought to investigate the more complex interplay between threatening objects and their spatial context. In real-world scenarios, threat perception is not determined by isolated features but emerges from their confluence. For instance, a tiger in a jungle evokes a potent threat response, whereas the same tiger behind zoo glass inspires little fear, illustrating that threat is perceived only when a threatening object appears in a contextually relevant location.

Therefore, Experiment 2a tested whether object-context interplay shapes the dynamics of attentional suppression. The primary hypothesis was that participants would develop learned suppression for high-probability locations, and the threat-associated salience would modulate the learned suppression. Specifically, if participants acquired suppression, we expected shorter response times (RTs) for the high-probability locations than the low-probability locations. Furthermore, if threat-associated salience potentiate learned suppression, we expected shorter RTs in the threat-associated salience objects presented at high-probability location than the non-threated-associated location.

## Materials and Methods

### Participants

The sample size calculation method and participant recruitment criteria were identical to those used in Experiment 1a. Forty-one participants were recruited for Experiment 2a. Behavioral data preprocessing followed the same protocol as in Experiment 1a. Following preprocessing, two participants were excluded due to a trial exclusion rate exceeding 30%. Consequently, a total of 39 participants (33 female; age range 19–25 years) were included in the final analysis.

### Stimuli and Apparatus

The stimuli and apparatus were largely identical to those used in Experiment 1a, with two key modifications. First, two distractor categories were presented, one associated with electric shocks (the threat-associated distractor, CsP) and one not associated with shocks (the non-threat-associated distractor, CsM). The assignment of the two categories to specific color was counterbalanced across participants. Second, the administration of shocks was contingent upon a specific combination of color and location. A shock was triggered only when the CsP appeared at a specific high-probability location; it was never triggered a shock when the same distractor appeared at another high-probability location. This design created three distinct distractor location conditions (see Figure 7B): the threat-associated distractor high-probability location (CsPH), the non-threat-associated high-probability location (CsMH) and low-probability distractor location (LPL).

**Figure 7.**
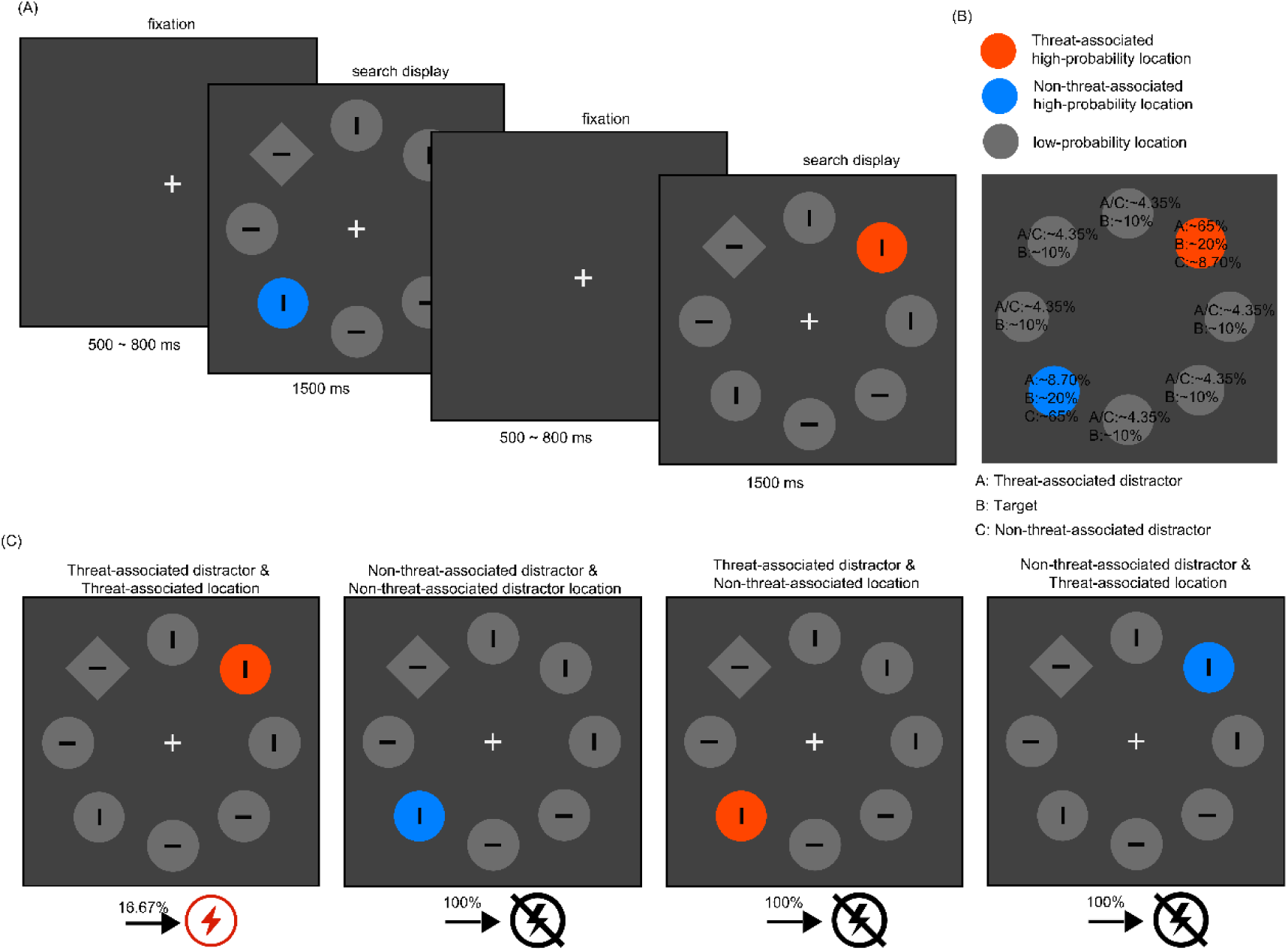
Tasks and Design for Experiment 2a *Note.* Panel A: Trial sequence of the visual search task. The task was to report the orientation of the bar inside the diamond. In the distractor-present condition, a color singleton replaces one of the non-target circles. Panel B: Example for the spatial and salience regularities of the distractor. The two high-probability distractor location are shown in orange and blue, while the low-probability location are shown in gray. Percentages at each location represent the probabilities of each distractor category or target to present in a given location. Panel C: Overview of the threat conditioning trial. The trial includes two distinct conditions: when the threat-associated distractor presents at the threat-associated high-probability location, approximately 16.67% of the trials are paired with an electric shock; in contrast, when threat-associated distractor presents at non-threat-associated high-probability location or any of distractor location, it is never paired with a shock. See the online article for the color version of this figure.

### Experimental Tasks

**Shock work-up task**: identical to that in Experiment 1a.

**Visual search Task**: The visual search task was generally consistent with that in Experiment 1a (see Figure 7A), with the following differences: Each block consisted of 124 trials: 92 distractor-present trials and 32 distractor-absent trials. In the distractor-present condition, six types of trials were designed to disentangle the effects of threat relevance and spatial probability. These included: (1) the threat-associated distractor presented at threat-associated high-probability location (CsP-CsPH, threat congruence, *trial number* = 30); (2) the threat-associated distractor presented at the non-threat-associated high-probability location (CsP-CsMH, threat incongruence, *trial number* = 4); (3) the threat-associated distractor presented at a low-probability location (CsP-LPL, *trial number* = 12, presented twice at each of 6 locations); (4) the non-threat-associated distractor presented at non-threat-associated high-probability location (CsM-CsMH, non-threat congruence, *trial number* = 30); (5) the non-threat-associated distractor presented at the threat-associated high-probability location (CsM-CsPH, non-threat incongruence, *trial number* = 4); and (6) the non-threat-associated distractor presented at a low-probability location (CsM-LPL, *trial number* = 12, presented twice at each of 6 locations). Notably, individuals are only at risk of being shocked when the CsP presents in CsPH. Specifically, five trials were randomly selected from 30 CsP-CsPH trials, and participants received an electric shock, corresponding to a probability of 16.67% (see Figure 7C). The shock was delivered at stimulus onset and was independent of the participants’ response. To prevent potential confounding effects, trials involving shocks were excluded from the analysis. In distractor-absent condition, target presented equally across the eight locations, with four times at each location.

**Implicit learning assessment**: identical to that in Experiment 1a.

## Data Analysis

The data analysis was largely identical to that used in Experiment 1a, after preprocessing, the proportion of trials removed ranged from 9.38% to 26.11%, with a mean of 16.64 ± 0.56%. The only difference was that in distractor-present condition, location probability and distractor category (CS) were included as fixed factors included as fixed effects with subject and target-distractor distance (TDD) as random effects (model: ACC/RT ∼ LocP + (1 | subject) + (1 | TDD)).

## Results

### Distractor-absent condition

As shown in Figure 8A, accuracy analysis revealed the main effect of distractor location was not significant (χ²_(2)_ = 0.61, *p* = 0.74).

**Figure 8.**
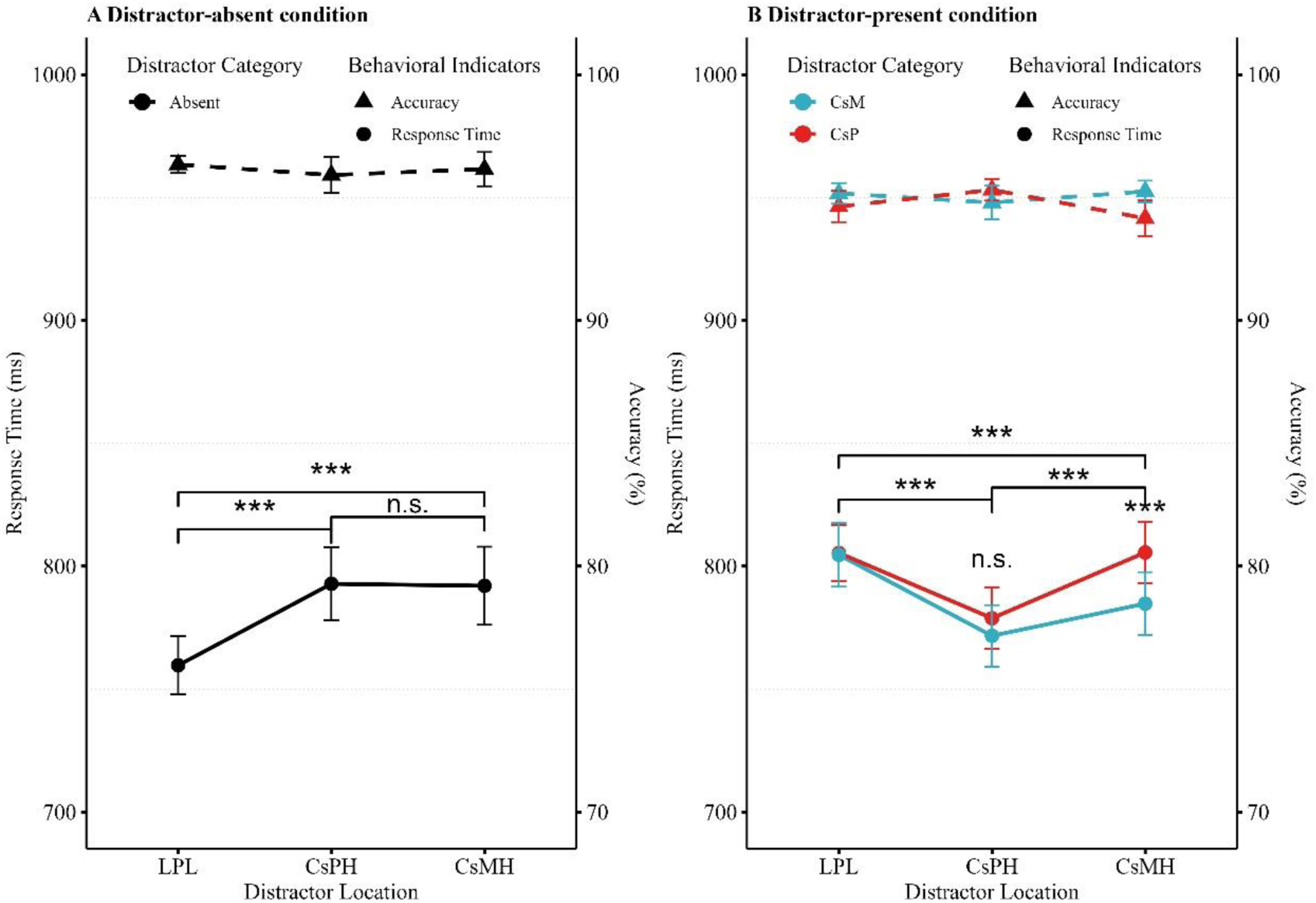
The results of search performance to target across different conditions in Experiment 2a. *Note.* Behavioral results for visual search in the distractor-absent condition (A) and distractor-present condition (B). The left Y-axis shows response time, represented by solid circles, and the right Y-axis shows accuracy, represented by solid triangles. All error bars represent the within-subject standard error of the mean. LPL: low-probability distractor locations; CsPH: threat-associated high-probability location; CsMH: non-threat-associated high-probability location. CsP: threat-associated distractor; CsM: non-threat-associated distractor. *** indicates *p* < 0.001, ** indicates *p* < 0.01, * indicates *p* < 0.05, and n.s. indicates not significant (*p* > 0.05). See the online article for the color version of this figure.

Response time analysis revealed a significant main effect of distractor location, *χ*²_(2)_ = 65.66, *p* < 0.001. Compared to low-probability location (LPL, 757.30 ± 11.76 ms), responses were slower at both threat-associated high-probability location (CsPH, 789.71 ± 16.08 ms, *b* = −31.23, *SE* = 5.13, *t*_(8,898)_ = 6.09, *p* < 0.001, 95% CI [21.18, 41.28]; *β* = 0.06, 95% CI [0.04, 0.08]) and non-threat-associated high-probability location (CsMH, 798.14 ± 15.55 ms, *b* = 31.51, *SE* = 5.12, *t*_(8,898)_ = 6.16, *p* < 0.001, 95% CI [21.48, 41.54]; *β* = 0.06, 95% CI [0.04, 0.08]). However, no significant difference was found between the two high-probability locations (*b* = 0.28, *SE* = 6.71, *t*_(8,898)_ = 0.04, *p* = 1.00, 95% CI [−12.87, 13.44]; *β* = 0.001, 95% CI [−0.02, 0.03]).

### Distractor-present condition

As shown in Figure 8B, accuracy analysis revealed neither significant main effects of distractor location(*χ*²_(2)_ = 2.21, *p* = 0.33), or distractor category (*χ*²_(1)_ = 1.51, *p* = 0.22), nor a significant interaction between them (*χ*²_(2)_ = 3.31, *p* = 0.19).

Response time analysis revealed a significant main effect of distractor location, (*χ*²_(2)_ = 65.66, *p* < 0.001). Compared to low-probability location, responses were faster both at threat-associated high-probability location (*b* = 29.94, *SE* = 3.39, *z* = 8.83, *p* < 0.001), and non-threat-associated high-probability location (*b* = 9.84, *SE* = 3.37, *z* = 2.918, *p* = 0.011). Additionally, responses were faster at threat-associated high-probability location compared to non-threat-associated high-probability location (*b* = −20.11, *SE* = 3.81, *z* = −5.28, *p* < 0.001). There was also a significant main effect of distractor category (*χ*²_(1)_ = 8.64, *p* = 0.003). Responses were slower when threat-associated distractors appeared than when non-threat-associated distractors appeared (*b* = 9.97, *SE* = 2.88, *z* = 3.46, *p* = 0.001). Furthermore, the interaction between distractor location probability and distractor category was significant (*χ*²_(2)_ = 9.71, *p* = 0.008). A post-hoc analysis showed that, compared to threat-associated distractor appeared at threat-associated high-probability location (CsP-CsPH, 778.82 ± 12.47 ms), responses were slower at both low-probability location (CsP-LPL, 805.38 ± 11.50 ms, *b* = 26.63, *SE* = 3.52, *z* = 7.57, *p* < 0.001) and non-threat-associated high-probability location (CsP-CsMH, 805.57 ± 12.50 ms, *b* = −27.28, *SE* = 5.42, *z* = −5.03, *p* < 0.001). In addition, compared to non-threat-associated distractor appeared at low-probability location (CsM-LPL, 804.62 ± 12.98 ms), responses were faster at both threat-associated high-probability location (CsM-CsPH, 771.64 ± 12.52 ms, *b* = 33.258, *SE* = 5.80, *z* = 5.73, *p* < 0.001) and non-threat-associated high-probability location (CsM-CsMH, 784.76 ± 12.81 ms, *b* = 20.32, *SE* = 3.42, *z* = 5.94, *p* < 0.001). Moreover, a slower response was observed when threat-associated distractor appeared at non-threat-associated high-probability location (CsP-CsMH) than non-threat-associated distractor appeared at non-threat-associated high-probability location (CsM-CsMH, *b* = 20.32, *SE* = 3.42, *z* = 5.94, *p* < 0.001).

## Awareness of statistical regularities

Eight of the 39 participants reported that they did not notice whether the distractor appeared more or less frequently in a specific location. These participants gave an average confidence rating of 3.25 ± 0.53 on a 7-point scale. The remaining 31 participants reported noticing a spatial imbalance and gave an average confidence rating of 3.71 ± 0.33. However, none of these participants correctly identified the high- or low-probability distractor locations as defined in the experiment. These results suggest that the learned suppression effect was driven by implicit awareness.

## Discussion

Experiment 2a replicated the robust learned suppression effect at high-probability locations, consistent with the findings of Experiments 1a and 1b. Under congruent conditions(i.e., CsP-CsPH and CsM-CsMH), distractor presence facilitated responses, indicating effective suppression at expected locations. However, an intriguing asymmetry emerge under incongruent conditions. Response times were significantly slower when a threat-associated distractor appeared at non-threat-associated high-probability location (i.e., CsP–CsMH), but not when a non-threat-associated distractor appeared at a threat-associated high-probability location (i.e., CsM–CsPH). This asymmetry suggests that a violation of learned object-location contingencies— specifically, the appearance of a threat distractor in a "safe" context—is treated as a salient statistical irregularity. This irregularity enhances the distractor’s perceptual salience, granting it pop-out properties that override the learned suppression and thereby capture attention.

While these trial-averaged results demonstrate how object-location congruence modulates suppression, they obscure the underlying temporal dynamics. It remains unclear whether the observed costs and benefits arise from differences in rapid reactive suppression, reduced attentional limbo, or earlier proactive suppression. To delineate these temporal profiles, Experiment 2b combined eye tracking with SMART analysis to reconstruct trial-by-trial trajectories of the first fixation. This approach allows us to precisely trace the time course of how threat associations and spatial context interact to shape attentional allocation from initial capture to sustained suppression.

## Experiment 2b

Building on the trial-averaged asymmetry observed in Experiment 2a, Experiment 2b leveraged a combination of eye tracking and SMART analysis to delineate the precise temporal dynamics through which threat associations and spatial context jointly shape attentional suppression. We hypothesized that learned suppression unfolds along a dynamic trajectory, transitioning from an initial reactive suppression to proactive suppression, often via an intermediate attentional limbo phase. Furthermore, we predicted that threat would critically modulate this progression. Specifically, we expected that the congruent pairing of a threat distractor with a threat-associated location (CsP-CsPH) would accelerate the shift from reactive to proactive suppression, minimizing time spent in attentional limbo. Conversely, we hypothesized that the incongruent appearance of a threat distractor in a safe location (CsP-CsMH) would disrupt this trajectory, resulting in prolonged reactive capture and a delayed development of proactive suppression.

## Materials and Methods

### Participants

The sample size calculation method and participant recruitment criteria were identical to those used in Experiment 1a. Forty-nine participants were recruited for Experiment 2b. Data preprocessing followed the same protocol as in Experiment 1b. Following preprocessing, six participants were excluded due to a trial exclusion rate exceeding 30%. Consequently, a total of 43 participants (38 female; age range 18-23 years) were included in the final analysis.

### Stimuli, Apparatus, Procedure

The stimuli and apparatus were identical to those used in Experiment 2a.

### Experimental Tasks

**Shock work-up task**: identical to that in Experiment 1a.

**Visual search Task**: The task was identical to those used in Experiment 2a, and the eye-tracking data recording procedure was followed the same protocol as in Experiment 1b.

**Implicit learning assessment**: identical to that in Experiment 1a.

## Data Analysis

The behavioral data analysis followed the same procedure as in Experiment 2a. After preprocessing, the proportion of trials removed ranged from 9.38% to 26.11%, with a mean of 16.64 ± 0.56%. The eye-tracking data analysis was similar to that of Experiment 1b. For the distractor-present condition, location probability and distractor category (CS) were included as fixed factors, and the probability of first fixation landing on the distractor served as the dependent variable. Distractor location (LocP) and distractor category (CS) were included as fixed factors, with subject and target-distractor distance (TDD) as random effects(model: 𝑃_𝑓𝑖𝑥_𝐷_ ∼ LocP ∗ CS + (1 ∣ Subject) + (1 ∣ TDD)).

## Results

### Behavioral results

#### Distractor-absent condition

As shown in Figure 9A, accuracy analysis revealed a significant main effect of distractor location (*χ*²_(2)_ = 7.80, *p* = 0.02). Compared to low-probability location (LPL, 97.03 ± 0.30%), accuracy was lower at both threat-associated high-probability location (CsPH, 96.07 ± 0.62%, *b* = −0.29, *SE* = 0.15, *z* = −1.92, *p* = 0.055, 95% CI [−0.59, 0.01], *OR* = 0.746) and non-threat-associated high-probability location (CsMH, 95.85 ± 0.84 %, *b* = −0.35, *SE* = 0.15, *z* = −2.34, *p* = 0.02, 95% CI [−0.64, −0.06], *OR* = 0.71). However, no significant difference was observed between the two high-probability locations (*b* = −0.06, *SE* = 0.19, *z* = −0.30, *p* = 0.77, 95% CI [−0.44, 0.32], *OR* = 0.95).

**Figure 9.**
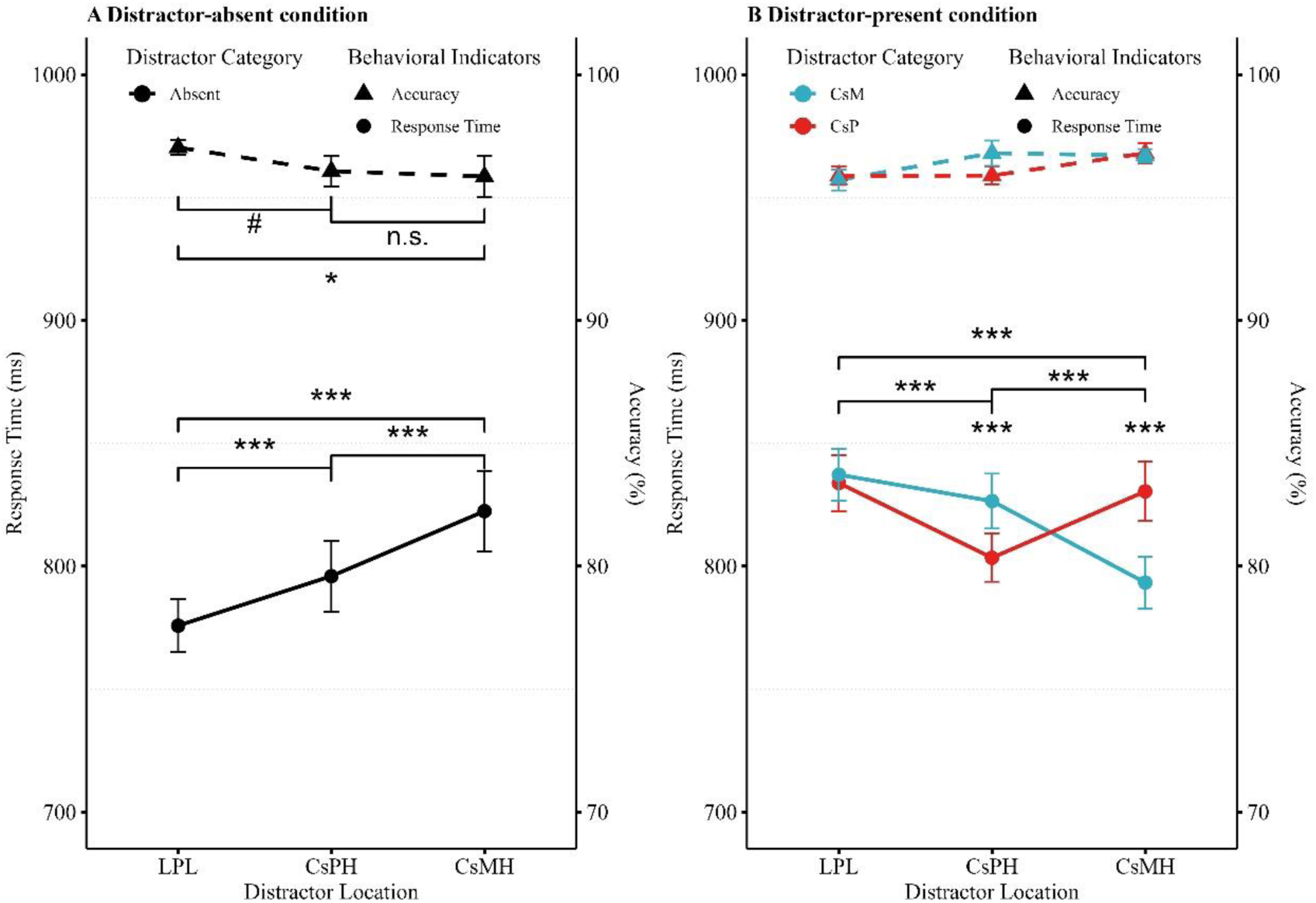
The results of search performance to target and first fixation landing probability across different conditions in Experiment 2b. *Note.* Behavioral results for visual search in the distractor-absent condition (A) and distractor-present condition (B). The left Y-axis shows response time, represented by solid circles, and the right Y-axis shows accuracy, represented by solid triangles. All error bars represent the within-subject standard error of the mean. LPL: low-probability distractor locations; CsPH: threat-associated high-probability location; CsMH: non-threat-associated high-probability location. CsP: threat-associated distractor; CsM: non-threat-associated distractor. *** indicates *p* < 0.001, ** indicates *p* < 0.01, * indicates *p* < 0.05, and n.s. indicates not significant (*p* > 0.05). See the online article for the color version of this figure.

Response time analysis revealed a significant main effect of distractor location (*χ*²_(2)_ = 100.38, *p* < 0.001). Compared to low-probability location (LPL, 775.82 ± 10.78 ms), responses were faster at both the threat-associated high-probability location (CsPH, 795.96 ± 14.41 ms, *b* = 19.865, *SE* = 4.712, *t*_(10,080)_ = 4.22, *p* < 0.001, 95% CI [10.628, 29.102]; *β* = 0.038, 95% CI [0.021, 0.056]) and non-threat-associated high-probability location (CsMH, 822.38 ± 16.35 ms, *b* = 45.147, *SE* = 4.705, *t*_(10,080)_ = 9.60, *p* < 0.001, 95% CI [35.925, 54.369]; *β* = 0.088, 95% CI [0.070, 0.105]). Responses were also faster at threat-associated high-probability location compared to the non-threat-associated high-probability location (*b* = 25.282, *SE* = 6.167, *t*_(10,080)_ = 4.10, *p* < 0.001, 95% CI [13.194, 37.371]; *β* = 0.049, 95% CI [0.026, 0.070]).

#### Distractor-present condition

As presented in Figure 9B, accuracy analysis revealed significant main effects of distractor location (*χ*²_(2)_ = 9.26, *p* = 0.01). Compared to low-probability location, accuracy was higher at non-threat-associated high-probability location (*b* = −0.27, *SE* = 0.10, *z* = −2.76, *p* = 0.018). However, no significant main effect of distractor category (*χ*²_(1)_ = 0.18, *p* = 0.67) or interaction with distractor location (*χ*²_(2)_ = 2.79, *p* = 0.25) was found.

Response time analysis revealed a significant main effect of distractor location, *χ*²_(2)_ = 222.73, *p* < 0.001. Compared to low-probability location, responses were faster at both threat-associated high-probability location (*b* = 20.37, *SE* = 3.19, *z* = 6.395, *p* < 0.001), and non-threat-associated high-probability location (*b* = 23.70, *SE* = 3.15, *z* = 7.527, *p* < 0.001). However, the main effect of distractor category was not significant, (*χ*²_(1)_ = 1.20, *p* = 0.27). The interaction between distractor location and distractor category was significant(*χ*²_(2)_ = 76.32, *p* < 0.001). A post-hoc analysis showed that, compared to threat-associated distractor appeared at the threat-associated high-probability location (CsP-CsPH, 803.42 ± 9.74 ms), response were slower at both low-probability location (833.84 ± 11.43 ms, *b* = 30.02, *SE* = 3.32, *z* = 9.05, *p* < 0.001) and non-threat-associated high-probability location (CsP-CsMH, 830.43 ± 12.08 ms, *b* = −26.76, *SE* = 5.04, *z* = −5.309, *p* < 0.001). Additionally, compared to non-threat-associated distractor at non-threat-associated high-probability location (CsM-CsMH, 793.33 ± 10.53 ms), responses were slower at both low-probability location (CsM-LPL, 837.20 ± 10.55 ms, *b* = 44.15, *SE* = 3.23, *z* =13.69, *p* < 0.001) or threat-associated high-probability location (CsM-CsPH, 826.48 ± 11.17 ms, *b* = 33.42, *SE* = 5.02, *z* = 6.66, *p* < 0.001). Critically, responses were slower when non-threat-associated distractor appeared at threat-associated high-probability location (CsM-CsPH) versus threat-associated distractor appeared at threat-associated high-probability location (CsP-CsPH, *b* = −22.57, *SE* = 5.08, *z* = −4.445, *p* < 0.001). Similarly, responses were slower for threat-associated distractor appeared at the non-threat-associated high-probability location (CsP-CsMH) than non-threat-associated distractor appeared at non-threat-associated high-probability location (CsM-CsMH, *b* = 37.61, *SE* = 4.98, *z* = 7.549, *p* < 0.001).

## Results of First Fixation Landing Probability

### Distractor-absent condition

As shown in Figure 10A, analysis of the first fixation landing on the target revealed a significant main effect of distractor location (*χ*²_(2)_ = 34.09, *p* < 0.001). Compared to low-probability distractor location (LPL, 41.23 ± 2.14%), the probability was lower at both threat-associated distractor high-probability location (CsPH, 35.51 ± 3.04%, *b* = −0.26, *SE* = 0.07, *z* = −3.89, *p* < 0.001, 95% CI [−0.39, −0.13], *OR* = 0.774) and non-threat-associated high-probability location (CsMH, 33.96 ± 3.25%, *b* = −0.32, *SE* = 0.07, *z* = −4.84, *p* < 0.001, 95% CI [−0.45, −0.19], *OR* = 0.73). However, no significant difference was observed between the two high-probability locations (*b* = −0.06, *SE* = 0.09, *z* = −0.74, *p* = 0.46, 95% CI [−0.24, 0.11], *OR* = 0.94).

**Figure 10.**
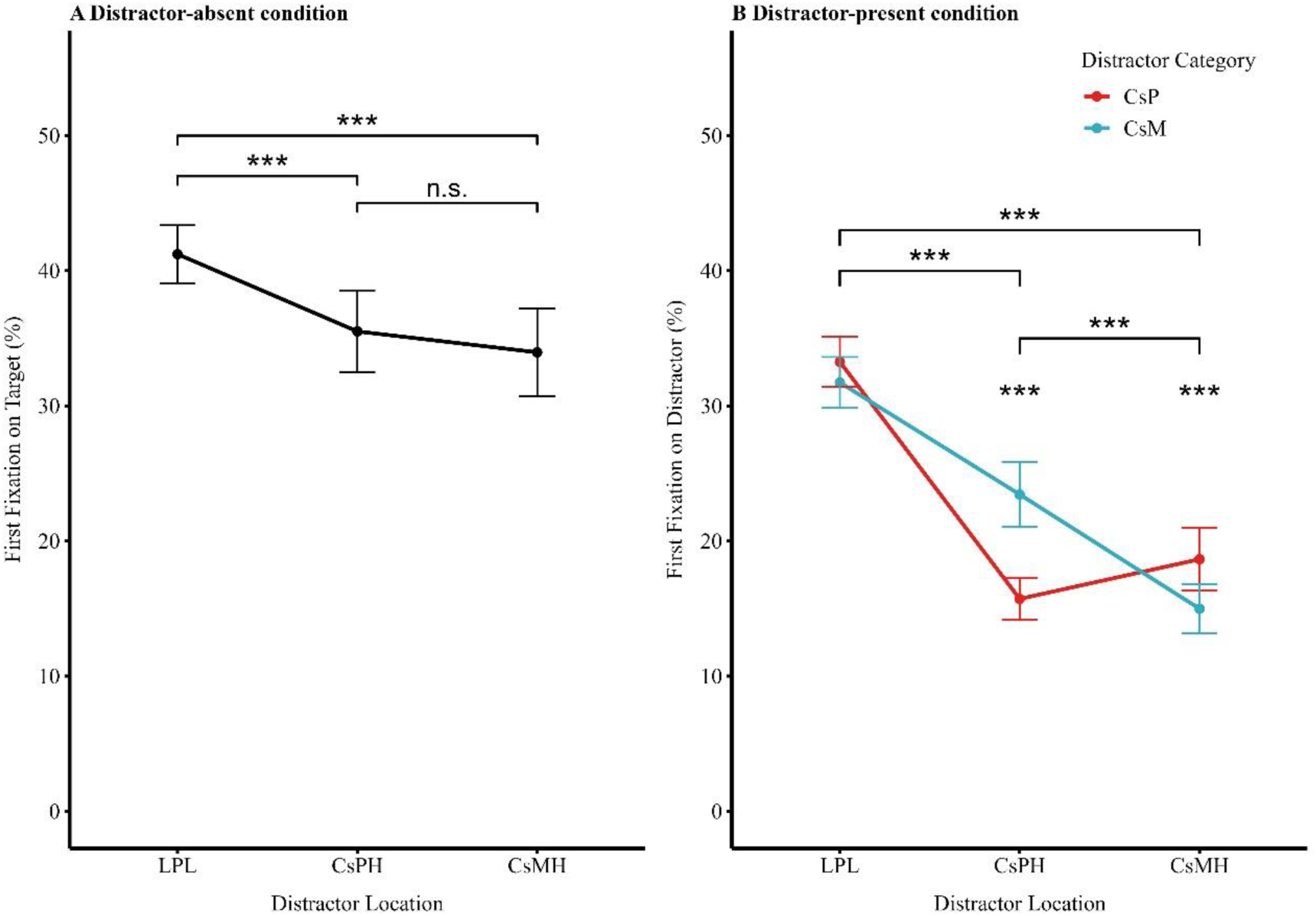
The results of first fixation landing probability across different conditions in Experiment 2b. *Note.* Panel A: The probability of the first fixation landing on the target when it appeared at a low-probability location, a threat-associated high-probability location, or a non-threat-associated high-probability location in the distractor-absent condition. Panel B: The probability of the first fixation landing on distractor when the distractor appeared at a low-probability location, a threat-associated high-probability location, or a non-threat-associated high-probability location in the distractor-present condition. All error bars represent the within-subject standard error of the mean. LPL: low-probability distractor locations; CsPH: threat-associated high-probability location; CsMH: non-threat-associated high-probability location. CsP: threat-associated distractor; CsM: non-threat-associated distractor. *** indicates *p* < 0.001, ** indicates *p* < 0.01, * indicates *p* < 0.05, and n.s. indicates not significant (*p* > 0.05). See the online article for the color version of this figure.

### Distractor-present condition

As shown in Figure 10B, analysis of first fixation landing on the distractor revealed a significant main effect of distractor location probability (*χ*²_(2)_ = 910.08, *p* < 0.001). Compared to low-probability location (32.50 ± 1.69 %), the probability was lower at both threat-associated high-probability location (16.8 ± 1.57%, *b* = 0.74, *SE* = 0.05, *z* = 16.43, *p* < 0.001) and non-threat-associated high-probability location (15.40 ± 1.78%, *b* = 0.93, *SE* = 0.05, *z* = 19.73, *p* < 0.001). Critically, a significant difference emerged between the threat-associated and the non-threat-associated high-probability location (*b* = 0.18, *SE* = 0.05, *z* = 3.33, *p* = 0.003). However, the main effect of the distractor category was not significant (*χ*²_(1)_ = 0.46, *p* = 0.50). The interaction between distractor location probability and distractor category was significant (*χ*²_(2)_ = 59.74, *p* < 0.001). A post-hoc analysis showed that, compared to threat-associated distractor appeared at the low-probability location (CsP-LPL, 33.26 ± 1.87%), the probability was lower at both threat-associated high-probability location (CsP-CsPH, 15.71 ± 1.54%, *b* = 1.03, *SE* = 0.05, *z* = 21.60, *p* < 0.001) and non-threat-associated high-probability location (CsP-CsMH, 18.65 ± 2.32%, *b* = 0.81, *SE* = 0.08, *z* = 10.03, *p* < 0.001). Additionally, compared to non-threat-associated distractor appeared at the low-probability location (CsM-LPL, 31.73 ± 1.89%), the probability was lower at both non-threat-associated high-probability location (CsM-CsMH,14.99 ± 1.82%, *b* = 1.04, *SE* = 0.05, *z* = 22.22, *p* < 0.001) and threat-associated high-probability location (CsM-CsPH, 23.44 ± 2.39%, *b* = 0.46, *SE* = 0.08, *z* = 5.98, *p* < 0.001). Moreover, the probability was lower when the threat-associated distractor appeared at the threat-associated high-probability location (CsP-CsPH) than non-threat-associated distractor appeared at the threat-associated high-probability location (CsM-CsPH, *b* = −0.50, *SE* = 0.07, *z* = −6.73, *p* < 0.001). Also, the probability was lower when the non-threat-associated distractor appeared at the non-threat-associated high-probability location (CsM-CsMH) than when the threat-associated distractor appeared at the non-threat-associated high-probability location (CsP-CsMH, *b* = 0.29, *SE* = 0.08, *z* = 3.64, *p* < 0.001).

## SMART Results for First Fixation Landing Probability

For the threat-associated distractors, at the low-probability locations (CsP-LPL, see Figure 11A), first fixations landed more frequently on distractors than targets during Trial 1-243 and Trial 261-376 (*ps* < 0.05). No significant difference emerged during Trial 244-260 and Trials after 377 (*ps* > 0.05). At threat-associated high-probability location (CsP-CsPH, see Figure 11B), first fixation landed on distractor initially on Trial 1-60 (*p* < 0.05), followed by a null phase where no significant difference emerged between landing on target and distractor on Trial 61-143 (*p* > 0.05), from Trial 144 onward, first fixation landed more frequently on target than distractor (*p* < 0.05). At non-threat-associated high-probability location (CsP-CsMH, see Figure 11C), no significant difference of first fixation landing was observed from Trial 1-251, Trial 320-362, Trial 389-563 (*ps* > 0.05). However, the probability of fixation on the target was higher than on the distractor during Trial 252-319, Trial 363-388, and Trial 564 onward (*ps* < 0.05).

**Figure 11.**
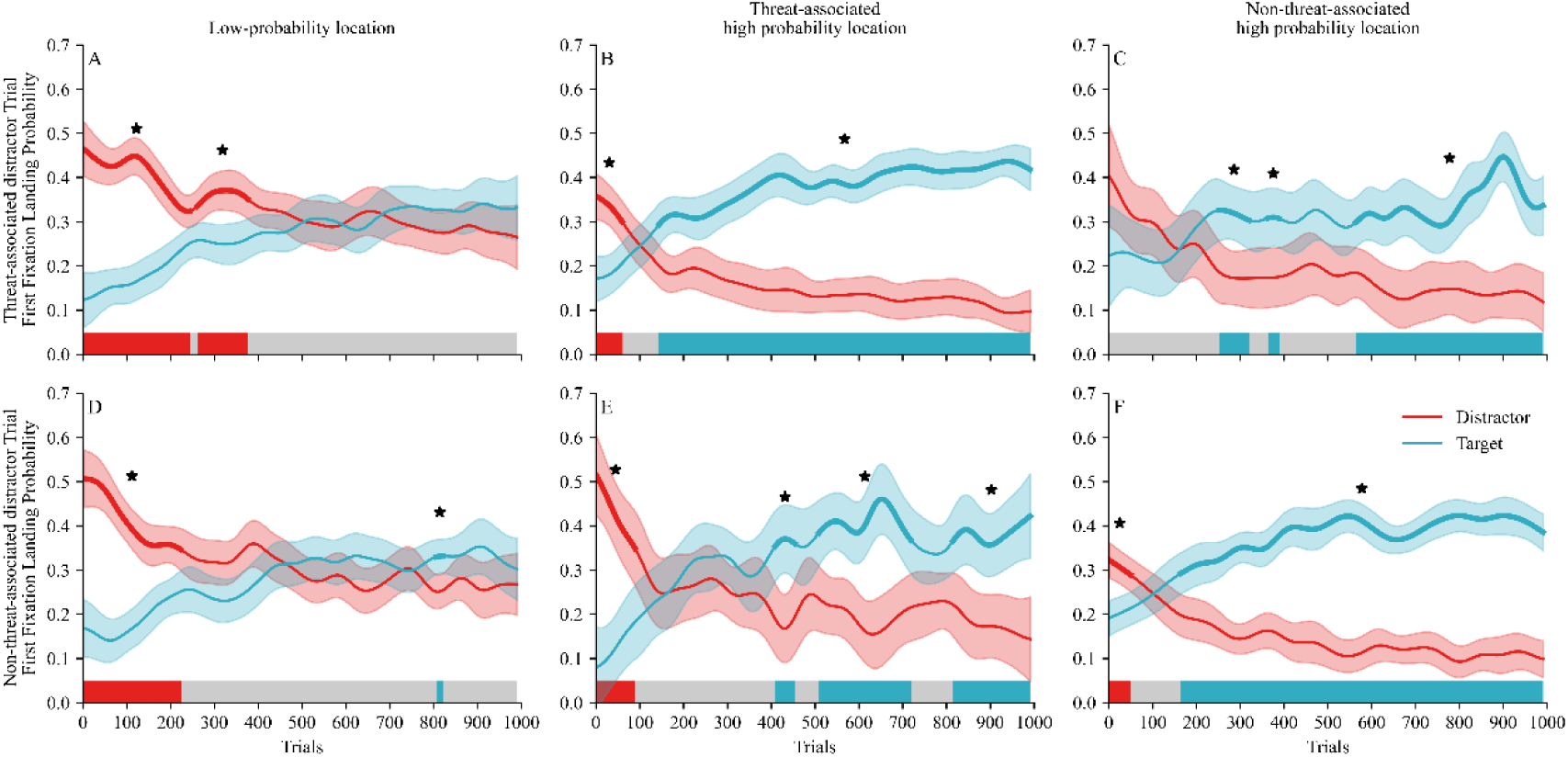
Difference functions reflecting the net distractor and target effects across trials in Experiments 2b *Note.* Panels A–C show data from threat-associated distractor trials for a low-probability location (panel A), a threat-associated high-probability location (panel B), and a non-threat-associated high-probability location (panel C). Panels D–F show data from non-threat-associated distractor trials for the low-probability location (D), the threat-associated high-probability location (E), and the non-threat-associated high-probability location (F). The shaded areas denote 95% confidence intervals, and the bold lines indicate the time points at which the probability of the first fixation differs significantly between the distractor and the target. * indicates *p* < 0.05. On the x-axis, the red-shaded region represents reactive suppression, the gray region represents attentional limbo, and the blue region represents proactive suppression. See the online article for the color version of this figure.

For the non-threat-associated distractors (CsM), at the low-probability location (CsM-LPL, see Figure 11D), the probability of the first fixation landing on the distractor was higher than on the target during Trial 1-223 (*p* < 0.05). However, no significant difference was observed during Trial 224-806 and Trial 823 onward (*ps* > 0.05). The probability of first fixation landing on the target was higher than on the distractor during Trial 807-822 (*p* < 0.05). At the threat-associated high-probability location (CsM-CsPH, see Figure 11E), the probability of first fixation landing on the distractor was higher than on the target during Trial 1-89 (*p* < 0.05). However, no significant difference was observed during Trial 90-408, Trial 455-508 and Trial 720-814 (*ps* > 0.05). The probability for the first fixation landing on the targets was higher than on distractors during Trial 409-454, Trial 509-719 and Trial 815 onward (*ps* < 0.05). At non-threat-associated high-probability location (CsM-CsMH, see Figure 11F), the probability of first fixation landing on the distractor was higher than on target during Trial 1-49 (*p* < 0.05). However, no significant difference occurred during Trial 50-163 (*p* > 0.05). From Trial 164 onward, the probability of first fixation landing on the target was higher than on the distractor (*p* < 0.05).

## Awareness of statistical regularities

Thirteen of the 43 participants reported that they did not notice whether the distractor appeared more or less frequently in a specific location. These participants gave an average confidence rating of 3.54 ± 0.35 on a 7-point scale. The remaining 31 participants reported noticing a spatial imbalance and gave an average confidence rating of 3.60 ± 0.30. However, none of these participants correctly identified the high-or low-probability distractor locations as defined in the experiment. These results suggest that the learned suppression effect was driven by implicit awareness.

## Discussion

Experiment 2b replicated the trial-averaged results of Experiment 2a, confirming robust learned distractor suppression at high-probability locations that critically depended on object–location congruence. Response was significantly faster when distractor features matched high-probability locations (i.e., CsP–CsPH, CsM–CsMH) than at low-probability locations. Conversely, incongruent conditions (i.e., CsM–CsPH, CsP–CsMH) elicited slower responses relative to congruent conditions. These findings underscore the pivotal role of feature–location congruence in shaping learned suppression.

SMART analyses of first-fixation landing probabilities reveal that learned distractor suppression operates through a dynamic attentional control states that evolve across learning phase. Critically, this evolution is modulated by both spatial probability and motivational salience in distinct stages. At low-probability locations, suppression transitions from reactive suppression to attentional limbo state, Threat associations prolong the reactive phase (capture persistence), delaying entry into limbo states. At high-probability locations, suppression progresses through three stages: from reactive suppression to attentional limbo and then to proactive suppression. This triphasic trajectory persists regardless of whether the distractor is threat-associated (CsP–CsPH), non-threat-associated (CsM–CsMH), or a non-threat-associated distractor presented at a threat-associated high-probability location (CsM–CsPH). Notably, the attentional limbo phase is significantly shortened for the CsP–CsPH condition, demonstrating threat-context congruence optimizes suppression efficiency. Finally, when threat-associated distractors appear at a safe high-probability locations (CsP–CsMH), suppression shifts directly from attentional limbo to proactive suppression. Critically, the replication of spatially specific suppression effects across experiments (2a vs. 2b) confirms that feature-location binding is a computational prerequisite for statistical learning of distractor suppression.

Together, these results suggest that learned distractor suppression is a dynamic process that changes over time. Suppression efficacy is determined by the congruence between spatial regularities and motivational salience.

## General Discussion

The current study investigated the mechanisms underlying learned distractor suppression and how they are modulated by motivational salience. Consistent with previous work^5–7^, we observed robust learned suppression effects in both trial-average behavioral measures and eye-tracking data. Critically, however, SMART-based analysis of first fixation probabilities revealed the temporal dynamics of learned suppression. Our findings demonstrate that learned distractor suppression is not a unified process but rather emerges from the dynamic integration of distinct reactive and proactive mechanisms within the attentional priority map over time. These results help resolve the ongoing debate regarding the mechanisms of learned distractor suppression and establish spatiotemporally distributed suppression as a core principle of attentional selection.

Across all four experiments, trial-averaged analyses consistently demonstrated robust learned distractor suppression, as evidenced by significantly faster responses when distractors appeared at high-probability locations but slower responses when targets appeared there. This behavioral pattern was corroborated at the oculomotor level by eye-tracking data: when distractors were present, first fixations were less likely to land on the distractor at high-probability locations; when distractors were absent, first fixations were less likely to land on targets at those locations. Thus, both behavioral and oculomotor measures converge to indicate that high-probability locations are subject to learned suppression, which was highly consistent with previous studies^29^.

## 1. Beyond physical salience: threat-related motivational salience enhances learned distractor suppression

Our study revealed that the mechanisms underlying learned distractor suppression differ markedly between high- and low-probability locations, highlighting the critical role of spatial expectation in shaping attentional suppression processes^66^. These results are consistent with those of Zhao et al. (2024), who found that distractors appeared at high-probability locations were associated with significantly larger pre-stimulus P_D_ component, suggesting a clear proactive (anticipatory) suppression mechanism. In contrast, distractors at low-probability locations elicited no such neural signatures and showed reduced suppression effects under high cognitive load, which is indicative of a greater reliance on reactive suppression. Building on this, our results further demonstrated that the suppression unfolds via distinct mechanisms over time. Specifically, suppression at low-probability locations primarily involved a transition from reactive suppression to attentional limbo state. Conversely, suppression at high-probability locations followed a more dynamic trajectory, progressing from reactive suppression, through attentional limbo, to proactive suppression. In some instances, a direct shift from attentional limbo to proactive suppression was observed. Together, these findings underscore the critical roles of spatial expectation and cognitive resource availability in shaping suppression trajectories. In essence, learning to anticipate distractor locations fosters not only a reconfiguration of attentional allocation but also the emergence of a resource-independent, proactively engaged suppression mechanism, which is fundamental to efficient attentional control.

Extending the prevalent view that physical salience enhances learned distractor suppression^9,65^, our findings provide evidence that motivational salience (e.g., threat) similarly potentiates this effect. In Experiment 1a, responses were faster when distractors appeared at the threat-associated high-probability locations than at the neutral high-probability locations. Moreover, Experiment 2 revealed that responses were faster under threat-congruent conditions than under threat-incongruent conditions. These findings demonstrate the synergistic interaction between learned spatial expectations and motivational experiences in shaping attentional control. Collectively, our results offer compelling evidence that individuals can learn to suppress motivationally salient distractors,including threat signals. These results align with growing evidence that threat-related stimuli—despite their inherent salience—can be effectively suppressed within predictable spatial contexts^60–62^. For instance, Nian et al. (2025) demonstrated that threat-history distractors were suppressed more efficiently than neutral ones at high-probability locations. Similarly, Theeuwes et al. (2025) showed that fear-conditioned stimuli (CS+) continued to be suppressed when appearing in previously suppressed locations, challenging the notion that threat-driven attentional capture is automatic and inflexible. Extending this, Theeuwes and van Moorselaar (2024) found that individuals with a spider phobia could suppress spider stimuli at high-probability locations, despite their heightened salience. By doing so, our work broadens the scope of distractor suppression theory beyond traditional visual salience to encompass motivational salience, thereby highlighting the important role of emotional and motivational factors in attentional learning.

Our findings further elucidate the critical role of threat in modulating the temporal dynamics of learned suppression. In Experiment 1b, suppression of distractors at a threat-associated high-probability location unfolded through a sequence from reactive suppression to attentional limbo and, ultimately, to proactive suppression. In contrast, at non-threatening high-probability location, suppression transitioned directly from attentional limbo to proactive suppression, bypassing the reactive stage. These patterns were validated in Experiment 2b, which showed that both threat-related and neutral distractors could evoke the full suppression trajectory (reactive-limbo-proactive) under specific spatial contingencies. Notably, the attentional limbo state was significantly shortened under threat conditions, implying that threat not only facilitates the early onset of suppression but also accelerates the transition to proactive control by compressing intermediate processing states.

Converging with our findings, this body of work supports a key theoretical insight: threat does not invariably override attentional suppression but can be dynamically integrated into learned control systems in a facilitative and adaptive manner. Crucially, our results extend the Signal Suppression 2.0 framework^67^ by not only confirming that threat-related distractors can be suppressed but also by mapping how threat modulates the temporal structure of this process. The accelerated transition from reactive to proactive suppression under threat suggests that threat may act as an internal cue that facilitates the updating of attentional priorities, potentially through mechanisms involving heightened arousal or enhanced motivational salience.

## 2. The primacy of learned distractor suppression over time: the *Temporal Dynamics of Attentional Suppression (TDAS)* hypothesis

The long-standing debate in learned distractor suppression research has centered on whether the underlying mechanism is primarily reactive or proactive. Mounting neuroscientific evidence, however, indicates that this dichotomy is overly simplistic, as neither mechanism alone can fully account for the complex, dynamic nature of learned attentional suppression ^24,66,68–71^. In response to this theoretical impasse, we propose the *Temporal Dynamics of Attentional Suppression (TDAS)* hypothesis. The core tenet of the TDAS hypothesis is that learned distractor suppression is not a unitary process but rather emerges from the dynamic integration of distinct suppression mechanisms within the priority map over time. The TDAS hypothesis framework delineate three temporally evolving state: (1) reactive suppression in which salient distractors dominate the priority map, triggering a stimulus-driven suppression; (2) attentional limbo in which target and distractor representations compete without a clear winner, resulting in a period of unresolved competition and belief updating; (3) proactive suppression in which through statistical learning, target representations gain dominace, enabling top-down, anticpatory suppression of the high-probability distractor location. Critically, this theoretical progression is directly and empirically supported by our SMART analyses of first-fixation landing probabilities, which revealed that suppression is not static but reflects dynamic transitions between the three states. Moreover, the precise trajectory is modulated by contextual factors: at low-probability locations, suppression typically shifted from reactive to attentional limbo; whereas at high-probability locations, it progressed sequentially through all three states. Or in some cases, transitioned directly from limbo to proactive suppression.

This TDAS hypothesis helps to resolve a fundamental theoretical tension by reconceptualizing suppression as a process of predictive priority remapping within a competitive integration architecture. In this view, the reactive state may reflect uncalibrated prediction errors in response to sudden salience. The limbo state represents a period of belief updating, where prior expectations are weighted against new evidence. The proactive state implements precision-weighted predictions, allowing for anticipatory allocation of attention. This framework is aligned with the empirical findings of Zhang et al. (2025), who using a forced-response method to track attentional priority over time, observed that attention fluctuated toward or away from salient distractors due to asynchronous target and distractor signals. Their computational modeling identified two complementary mechanisms: a slow suppression, where initial distractor dominance is gradually overridden by the target; and a fast suppression, which directly inhibits distractor influence^43^. These map neatly onto the TDAS framework, with the slow mechanism corresponding to the transition from reactive suppression to attentional limbo, and the fast mechanism facilitating the shift to proactive control.

Furthermore, the Signal Suppression 2.0 framework provides crucial cognitive and neurophysiological support for the TDAS model. It posits that learned suppression relies on the implicit learning of specific features and spatial locations, a process often initiated by an initial attentional capture of the distractor^67^. This account explains the counterintuitive finding that high-salience distractors are paradoxically easier to suppress and why proactive suppression typically emerges only after reactive capture has occurred. Thus, Signal Suppression 2.0 elucidates the learning pathway and mechanistic foundation that underpins the transition from reactive to proactive suppression described by the TDAS model.

The TDAS model integrates these diverse lines of evidence into a unified framework, establishing spatiotemporally distributed suppression as a core principle of attentional selection. It moves beyond the reactive-proactive dichotomy by explaining how the system dynamically adapts its suppression strategy based on temporal evolution and contextual demands. To advance this line of research, future studies could pursue several important directions. First, it is essential to investigate whether the observed temporal dynamics of suppression generalize to other classes of salient distractors, such as abrupt onsets or dynamic stimuli, which are typically more resistant to suppression. Examining these stimuli will help delineate the boundary conditions of generalizability of suppression flexibility. Second, employing time-resolved neuroimaging techniques—such as EEG measures of the P_D_ component or fMRI decoding of frontal eye field activity—would allow for a more precise characterization of the neural transitions between reactive and proactive suppression phases, as well as help disentangle the contributions of cortical versus subcortical systems. Third, future work should explore how individual differences—including working memory capacity, trait anxiety, and attentional control efficiency—modulate suppression flexibility, particularly under emotionally charged conditions that impose greater demands on cognitive control.

In a word, our findings underscore the temporal flexibility of attentional suppression in response to motivationally salient stimuli, providing novel insights into the dynamic interplay between statistical learning and emotional salience.

## Author Contributions

Jingqing Nian served as lead for data curation, formal analysis, and validation and contributed equally to conceptualization, writing–original draft, and writing–review and editing. Di Zhang served as lead for conceptualization, and review and editing. Yu Zhang served as lead for conceptualization, supervision, and writing–review and editing. Yu Luo served as lead for conceptualization, funding acquisition, supervision, writing–original draft, and writing–review and editing.

## Acknowledgements

This research was supported by the Basic Research Program of Guizhou Province (Qiankehe Jichu -ZK[2023] General-276) and the National Natural Science Foundation of China (32260209).

